# Human- and Rodent-derived Extracellular Vesicles Mediate the Spread of Pathology in MSA-like Models

**DOI:** 10.1101/2025.05.12.653337

**Authors:** Maria Vetsi, Dimitra Dionysopoulou, Panagiota Mavroeidi, Fedra Arvanitaki, Grigoria Tsaka, Hele Matinopoulos Lopez, Marianna Giannopoulou, Sotirios Fortis, Anastasios Kriebardis, Stefan Becker, Milan Zachrdla, Maria-Sol Cima Omori, Ismini Kloukina, Ioanna Tremi, Sofia Havaki, Vassilis G. Gorgoulis, Nadia Stefanova, Poul H. Jensen, Markus Zweckstetter, Maria Xilouri

## Abstract

Multiple system atrophy (MSA) is characterized by the presence of protein-rich inclusions mainly within oligodendrocytes, comprised primarily by the neuronal protein αSynuclein and the oligodendroglial-specific phosphoprotein TPPP/p25α. Μature oligodendrocytes do not normally express detectable αSynuclein levels, suggesting that its oligodendroglial accumulation may arise from intercellular transfer, potentially via extracellular vesicles (EVs); however the precise role of oligodendroglial-derived EVs in MSA progression remains relatively understudied.

Herein, we characterized the cargo/features and pathogenic potential of EVs released by oligodendrocytes treated with human αSynuclein fibrils amplified from MSA or Parkinson’s disease patient brains (or human recombinant αSynuclein fibrils) and EVs isolated from murine and human MSA (or respective control) brains. Our findings reveal that both oligodendroglial cell- and brain-derived EVs harbor pathological αSynuclein and TPPP/p25α conformations, similar to those accumulating in human MSA brains. These EVs are readily taken up by both neurons and oligodendrocytes, driving αSynuclein propagation *in vitro*.

Importantly, inoculation of these MSA-like EVs in animal models induce robust pSer129-αSynuclein accumulation along the nigrostriatal axis, colocalizing with markers of mature oligodendrocytes and dopaminergic neurons. These findings underscore the pivotal role of oligodendroglial-derived EVs in pathology progression and neuronal-oligodendroglial communication, positioning them as promising targets for therapeutic strategies aimed at combating alpha-Synucleinopathies.

## Introduction

alpha-Synucleinopathies, including multiple system atrophy (MSA) and Parkinson’s disease (PD), are characterized by the pathological accumulation of the neuronal protein αSynuclein (αSyn) [1]. Although both diseases share this common hallmark, they exhibit distinct cellular and pathological profiles. Specifically, αSyn accumulates predominantly within the glial cytoplasmic inclusions (GCIs) found mainly in oligodendrocytes of MSA brains, which drive widespread demyelination and neuronal degeneration [2]. Conversely, in PD, αSyn accumulates predominantly in neurons in the form of Lewy bodies and neurites that contribute to progressive neurodegeneration, particularly within the dopaminergic system [3, 4]. These differences in cellular targeting are attributed to differential cell-type-specific mechanisms and conformational strains of αSyn, which drive distinct propagation dynamics and toxicity across these disorders [5].

Physiologically, αSyn is a presynaptic neuronal protein that regulates synaptic vesicle trafficking and neurotransmitter release. However, in disease states, it undergoes pathological misfolding, aggregation and spreading. Reported scenarios link αSyn spread to a prion-like process, involving the release of misfolded αSyn aggregates into the extracellular space, their uptake by neighboring cells and subsequent seeding of aggregation [6, 7]. The ectopic localization of αSyn selectively to oligodendrocytes adds further complexity to MSA pathogenesis. Notably, the interaction of αSyn with the tubulin polymerization-promoting protein p25α (TPPP/p25α), an oligodendrocyte-specific protein critical for microtubule stability, has been implicated in exacerbating αSyn aggregation and accumulation, converting p25α’s physiological role into a pathological driver of disease progression [8, 9].

The intercellular transmission of αSyn aggregates is a hallmark of alpha-synucleinopathies, with extracellular vesicles (EVs) playing a central role in this process. Their cargo reflects the physiological or pathological state of the originating cell, rendering them critical players in disease propagation. Under pathological conditions, they have been implicated as carriers of pathogenic αSyn across the brain [10]. Interestingly, the intravesicular environment promotes αSyn aggregation [11] and EV-associated αSyn exhibits enhanced internalization and toxicity in recipient cells compared to EV-free protein [12]. Furthermore, EVs isolated from the cerebrospinal fluid of PD patients can induce oligomerization of soluble αSyn in target cells, facilitating the prion-like spread of pathology [13].

Oligodendroglial-derived EVs are gaining attention in alpha-synucleinopathy research, particularly in MSA. Their reduced secretion in the blood of MSA patients as compared to PD or control subjects has been associated with pathological αSyn aggregation [14]. While neuronal-derived EVs have been extensively studied, the role of oligodendroglial EVs remains relatively underexplored, especially in the context of MSA as unique oligodendrogliopathy. This study addresses this gap by comprehensively characterizing EVs derived from oligodendroglial cells treated with human αSyn fibrils amplified from MSA and PD brains or recombinant αSyn fibrils, along with EVs isolated from murine and human MSA patient (or control) brains, employing a battery of *in vitro* and *in vivo* models. By analyzing EV content, recipient cell responses and downstream effects on cellular and network integrity, we aimed to elucidate the contribution of oligodendroglial EVs—and the EV-associated αSyn and TPPP/p25α cargo—to αSyn propagation in the context of MSA. Our results indicate that oligodendroglial-derived EVs operate as carriers of pathological αSyn and TPPP/p25α protein conformations, both *in vitro* and *in vivo*. Additionally, we highlight a role of EVs isolated from a MSA mouse model and human MSA brains in the spread of αSyn-related pathology in the living brain. Most importantly though, these results expound the importance of oligodendroglial-derived EVs for neuronal-oligodendroglial crosstalk, harboring thus a considerable potential for the development of therapeutic interventions against MSA.

## Materials and Methods

### Study design

The aim of our study was to comprehensively characterize EVs released from αSyn fibril-treated oligodendroglial cell lines, mouse and human brains (MSA and respective controls), in terms of content, recipient cell responses (primary cultures and animal models) and downstream effects on cellular and network integrity. No sample size computations were performed during the experimental design. For the *in vivo* intrastriatal EV injections, three-month-old male WT C57BL6/C3H (WT-αSyn) or C57BL6 PLP-hαSyn transgenic mice were utilized. One- and three-months post-EV injections were selected at endpoints to observe time-dependent alterations on αSyn-related pathological indices. Similarly, 48h and 7 days endpoints were chosen for the *in vitro* experiments. All presented data represent the mean of at least three biological replicates. Animal selection, cell counting and image acquisition were performed randomly, by independent investigators blinded to sample identity until final data curation using number coding. All experiments include the appropriate control group and all collection and analysis settings were similar between groups.

### Cell culture and treatments

OLN-93, an immortalized oligodendroglial cell line originating from primary Wistar rat brain glial cultures [15], was transfected with either a pcDNA3.1 zeo(−) human αSyn vector or a pcDNA3.1 zeo(−) human p25α vector to establish the OLN-AS7 and OLN-p25α cell lines, respectively [16]. All cells were cultured in Dulbecco’s Modified Eagle Medium (DMEM, D6429, Gibco, Invitrogen), supplemented with 10% Fetal Bovine Serum (FBS, 10270, Gibco, Invitrogen), 50 U/mL penicillin and 50 μg/mL streptomycin. To sustain OLN-AS7 and OLN-p25α cells, 50 μg/mL Zeocin (R25001, Thermo Fisher Scientific) was added to the growth medium.

### Primary cortical neuron cultures

Cortices from P0 to P3 mouse brains were dissected and incubated in a papain solution (30 U/mL, Worthington, LS003120) with 180 U/mL DNase I (Invitrogen, Cat. No.: 18047-019) for 30 min at 37°C. Following centrifugation (400 g, 5 min, RT) the neuron-containing pellet was collected and resuspended in Neurobasal medium (Gibco, Cat. No.: 21103-049) supplemented with 2% (w/v) B-27 supplement (Gibco, Cat. No.: 17504-044), 2 mM L-glutamine (Glutamax, Gibco, Cat.No.: 35050-061), 1% penicillin/streptomycin (Biowest, Cat. No.: L0022-100) and 0.5% FBS. Neurons were then plated onto Poly-D-Lysine-coated (PDL, Sigma, Cat. No.: P7405) glass coverslips and seven days later were treated with the respective EVs.

### Primary oligodendroglial cultures

Primary oligodendroglial progenitor cells (OPCs) were isolated from mixed glial cell cultures prepared from P0 to P3 mouse brains, as previously described [17]. Subsequently, the isolated OPCs were plated on PDL-coated glass coverslips and cultured in SATO medium (Bottenstein and Sato, 1979) supplemented with Insulin-Transferrin-Selenium solution (41400045, Gibco, Invitrogen), 1% penicillin/streptomycin and 1% horse serum (H1138; Sigma Aldrich) to promote differentiation. Seven days later, cultures were treated with EVs.

### Preparation of brain-derived αSyn fibrils of MSA- and PD-patients

MSA or PD patient-derived fibrils were generated, as previously described [18]. Specifically, the protein misfolding cyclic amplification technique was employed to amplify αSyn fibrils from brain extracts of patients pathologically confirmed with MSA or PD (MSA and PD fibrils, respectively). Patient demographics are detailed in **Table S1**. In all experiments, cells were incubated with 1 or 6 μg/mL MSA or PD fibrils (or PBS as control) for 48h or 8 days.

### Preparation of human αSyn Pre-Formed Fibrils (PFFs)

The conversion process of monomeric αSyn into αSyn PFFs was performed as in [17]. For all experiments, cells were incubated with either 1 or 6 μg/mL PFFs (or PBS as control) for 48h or 8 days.

### Cell-derived EV purification

Cells were cultured in 150 mm dishes (Sarstedt, Cat. No: 83.3903) using DMEM supplemented with 10% FBS and 1% penicillin/streptomycin. 48h before EV isolation, fibril-contained medium was removed, cells were extensively washed three times with PBS and EV-depleted medium (prepared as described in Supplementary Methods) was added to the cultures. At the end of incubation time, the medium was collected, centrifuged at 400 g for 5 min and then subjected to ultracentrifugation at 100.000 g for 2 h at 4°C to isolate the EV-containing pellet. Finally, the pellet was resuspended in 50 μL PBS and stored at 80°C until further analysis. For lysosomal pathway inhibition, cells were either incubated with EV-depleted medium for 48h as normally or for the last 16 h of this 48h period the EV-depleted medium was supplemented with 20 mM ammonium chloride (NH₄Cl, general lysosomal inhibitor) before cell lysis and EV isolation.

### Autopsy case material

Human post-mortem brain tissue was obtained from the Queen Square Brain Bank, UK. This study involved two confirmed MSA cases and two controls without any neurological disease (the demographics of which are shown in **Table S2**). Specifically, EVs were isolated from the cerebellar white matter, a brain region with severe GCI pathology in MSA. The Ethical Committee of BRFAA approved the protocols for the use of these brains.

### Brain-derived EV isolation and purification

Mouse and human brain EVs were isolated as previously described [19], with minor adjustments. Briefly, brain tissues from mice lacking endogenous αSyn expression (KO-αSyn), wild-type (WT-αSyn) mice and mice overexpressing human αSyn selectively in oligodendrocytes, under the PLP promoter (PLP-hαSyn) or frozen human brain samples were digested with 20 units/mL papain in MEM medium for 30 min at 37°C, followed by homogenization in cold MEM. The homogenate was then sequentially filtered through a 40 µm mesh filter, followed by 0.45 and 0.22 µm syringe filters. The filtrate was centrifuged at 300 g (15 min, 4°C), 2.000 g (10 min, 4°C) and 10.000 g (30 min, 4°C). The resulting supernatant was centrifuged at 100.000 g (70 min, 4°C) to pellet the EVs. The EV-containing pellet was diluted in cold PBS and centrifuged again at 100.000 g (70 min, 4°C). The washed EV pellet was resuspended in 2 mL of 0.95 M sucrose solution, loaded on a sucrose step gradient and centrifuged at 200.000 g (16 h, 4°C). The obtained fractions were diluted in cold PBS and centrifuged at 100.000 g (70 min, 4°C). The resulting EV-enriched pellets from the sucrose gradient were resuspended in 50 µl of cold PBS and stored at −80°C for further analysis.

### Nanoparticle tracking analysis (NTA) of EV preparations

Particle size distribution and concentration of cell- and brain-derived EV preparations were assessed using NTA on the Nanosight NS300 instrument (Malvern Instruments, Amesbury, UK). The NanoSight NS300 features a 532 nm laser (green), a high sensitivity sCMOS camera and a syringe pump. EV preparations were diluted in particle-free PBS (0.22 µm filtered) to obtain a concentration within the recommended measurement range of 1-10 × 10^8^ particles/mL, corresponding to 1:100 dilution of the initial sample concentration. Each sample was loaded into a 1 ml syringe attached to the pump. During each measurement, five 30-second videos were captured under standardized conditions: EV temperature set at 25°C and syringe speed maintained at 100 µL/sec. Videos were then analysed using NanoSight NTA 3.4 build 3.4.4 software (Copyright 2020, Malvern) following capture in script control mode. A total of 1500 frames were examined per sample.

### Transmission Electron Microscopy (EM)

#### EM of fibrils morphology

Isolated αSyn fibrils were negative strained according to [17] protocol and observed in a FEI Morgagni 268 Transmission Electron Microscope (Netherlands) operated at 80kV and equipped with the Olympus Morada digital camera (Germany).

#### EM of EVs morphology

EVs were isolated, fixed, stained and embedded according to [20] protocol. They were observed in a FEI Morgagni 268 Transmission Electron Microscope (Netherlands) operated at 80kV and equipped with the Olympus Morada digital camera (Germany).

#### EM-Immunogold labeling of EVs

Immunogold labeling of EVs was based on protocol [20], with modifications. Briefly, isolated EVs were fixed in 2% paraformaldehyde (PFA) and adsorbed on Nickel Formvar/carbon-coated EM grids (200 mesh). EVs were then permeabilized with 0.15% saponin in PBS, blocked with 5% Normal Goat Serum (NGS)/1% BSA in PBS and incubated with primary antibodies against human αSyn (1:100), rodent αSyn (1:100), TPPP/p25α (1:100) or pSer129-αSyn (1:200) for 45 min at RT. Following rinsing, EVs were incubated with UltraSmall Gold-conjugated antibodies (0.8nm gold particles) (1:80) for 45 min at RT, and then subjected to silver enhancement for 4 min (NanoProbes, Silver Enhancement kit). Afterwards, EVs were washed, fixed with 1% glutaraldehyde, contrasted with uranyl oxalate and finally embedded in a mixture of 4% uranyl acetate and 2% methylcellulose. EVs were observed using a Jeol JEM2100Plus Transmission Electron Microscope (Japan) operated at 120kV and photographed with the Gatan OneView CMOS camera.

### Subcellular fractionation and Western immunoblotting

Cellular pellets underwent lysis with buffers of increasing extraction strength, supplemented with protease (Roche, 11836170001) and phosphatase (Roche, 04406837001) inhibitors. Initially, cells were treated with 1% Triton X-100 buffer (150 mM NaCl, 50 mM Tris pH 7.6, 2 mM EDTA) and centrifuged at 13.000 g for 30 min at 4°C. The supernatant (Triton-soluble fraction) was collected, while the pellet after two PBS-washes was resuspended in a 1% SDS buffer (150 mM NaCl, 50 mM Tris pH 7.6, 2 mM EDTA), sonicated and centrifuged to obtain the SDS-soluble fraction. Subsequently, the remaining pellet was washed and dissolved in an 8 M Urea-5% SDS buffer, heated for 30 min at 45°C and centrifuged to yield the Urea-soluble fraction. For EV-associated lysates, 5-minute bath-sonication was performed. Then, samples of equal protein concentration were subjected for Western blot analysis. Each immunoreactive band’s intensity was quantified using the ImageJ software. Differences in protein expression levels were assessed following normalization of all measurements using loading controls (β-actin for intracellular proteins and alix, flottilin-1, TSG101 or CD9 for EV-associated proteins).

### Immunocytochemistry (ICC) and confocal microscopy

Following fixation with 4% PFA, OLN cells and primary cultures were blocked (10% NGS, 0.4% Triton in PBS), incubated overnight with primary antibodies (2% NGS, 0.1% Triton in PBS), as detailed in **Table S3**, followed by secondary antibody incubation (2% NGS, 0.1% Triton in PBS). Nuclei were stained using the dye 4΄,6-diamidine-2΄-phenylindole dihydrochloride (DAPI), whereas PKH26 dye labeled EVs, enabling their uptake verification. Images were captured using a Leica TCS SP5 confocal microscope equipped with dual (tandem) scanner. ImageJ (v2.0.0) software was employed to quantify relative protein levels expressed as area coverage (μm^2^), normalized to the total number of cells per field.

### Animals

Three-month-old male WT C57BL6/C3H (WT-αSyn), C57BL6/JOlaHsd αSyn null (KO-αSyn) (purchased from Harlan Laboratories), or C57BL6 PLP-hαSyn transgenic mice were housed in the Animal House Facility of the Biomedical Research Foundation of the Academy of Athens (Athens, Greece). Animals were kept in individually ventilated cages (6-8 animals per cage) under pathogen-free conditions with free access to food and water on a 12-hour light/dark cycle. All animal experiments are approved by the Institutional Animal Care and Use Committee of Biomedical Research Foundation of Athens (792305 and 1392883 licence numbers), according to the European Legal framework existing for the Protection of animals used for experimental and other scientific purposes (European Convention 123/ Council of Europe and Directive 63/2010 of the European Union), along with the current Guidelines of International Organizations such as the Association for the Assessment and Accreditation of Laboratory Animal Care International-AAALAC Int., and the Federation of European Laboratory Animal Science Associations (FELASA).

### Surgical procedures

For stereotactic EV injections into the right dorsal striatum, mice were anesthetized with isoflurane (Abbott, B506) and placed in a stereotactic frame (Kopf Instruments, Tujunga, CA, USA). This region was targeted using the following bregma-based coordinates: +0.5 mm anterior-posterior, −1.8 mm (and −2.2 mm in case of OLN-p25α-derived EVs) medial-lateral, −3.4 and −3.2 mm dorsal-ventral, according to the Mouse Brain Atlas [21]. Using a pulled glass capillary, connected to a Hamilton syringe with a 22-gauge needle, EV suspension (2 μL per injection site) was injected at a rate of 0.1 μL per 15 seconds unilaterally. The following preparations were used: *a)* 10 μg PKH26-labeled WT-αSyn-derived EVs (PLP-hαSyn mice were sacrificed 7 days post-injection), *b)* 16 μg WT-αSyn- and PLP-hαSyn-derived EVs (WT-αSyn mice were sacrificed one and three months post-injection), *c)* 10 μg human-derived EVs (PLP-hαSyn mice were sacrificed three months post-injection) and *d)* 13 μg MSA- or PBS-treated OLN-p25α-derived EVs (PLP-hαSyn mice were sacrificed three months post-injection). The needle remained in place for 5 more min before being slowly withdrawn.

### Immunohistochemistry (IHC)

Experimental animals were anaesthetized with isoflurane and then subjected to intracardial perfusion with ice-cold PBS, followed by ice-cold 4% PFA solution. Their brains were dissected, post-PFA fixed for 24 h and transferred to 15% and then 30% sucrose. Brains were rapidly frozen in isopentane at −45°C for 60 seconds, before storage at −80°C. Then they were sectioned coronally at 30 μm increments using a Leica cryostat at −25°C. Immunohistochemical staining was performed on free-floating sections. Briefly, the sections were blocked (5% NGS, 0.3% Triton-X 100) and incubated with primary and secondary antibodies detailed in the Immunocytochemistry and confocal microscopy section.

### Imaris imaging software analysis

pSer129-αSyn within TH-positive neurons in the SNpc was quantified using the Imaris software (version 10.0.0). Initially, a surface was created for green channel (TH) to delineate TH-positive neurons. Subsequently, the red channel (pSer129-αSyn) was masked based on the TH surface, setting the external pixel-values to zero. A new surface was then generated from the masked red channel to identify pSer129-αSyn^+^ signal within TH-positive neurons. Automated statistical analyses provided by Imaris were utilized to measure following parameters: TH volume, total number and volume of phosphorylated puncta, as well as overlapped volume ratio between TH and pSer129-αSyn surfaces.

### Statistical analysis

Data were statistically analysed with GraphPad Prism 5. Differences within or between groups were assessed by one-way and two-way ANalysis Of VAriance (ANOVA) followed by Tukey’s post-hoc test and Bonferroni’s correction, respectively. Results are displayed as the mean ± Standard Error (SE), with a p-value of < 0.05 regarded as statistically significant. Results are based on the analysis of at least three independent experiments.

## Results

### Addition of human patient-amplified (MSA or PD) or recombinant αSyn fibrils alters the amount of oligodendroglial-derived EVs and increases the EV-associated release of αSyn species

Our aim was to assess the potential involvement of EVs with oligodendroglial origin in the transmission of αSyn-related pathology, by characterizing oligodendroglial-derived EV preparations under baseline conditions and following administration of human patient-derived or recombinant αSyn fibrils (1 or 6 μg/mL for 48h or 8 days). To evaluate the differential impact of human αSyn fibril strains originated mainly from oligodendrocytes (MSA fibrils) or neurons (PD fibrils), we utilized the respective patient-derived fibrils. EM analysis indicated distinct morphological features between MSA and PD fibrils, while heterogeneity was also observed between the two MSA fibril types (Fig.S1A). Immunoblotting unraveled different protein patterns across MSA1, MSA2, PD fibril samples and αSyn PFFs that became more pronounced following incubation with proteinase K (Fig.S1B). MSA1 fibrils demonstrated greater resistance to proteinase K digestion compared to MSA2 and PD fibrils. This structural variability may correlate with the patients’ distinct pathological phenotypes from whom the fibrils originated (**Table S1**). Application of all fibril types [1 or 6 μg/ for 48h or 8 days (or PBS as control)] to oligodendroglial (OLN) cell lines resulted in increased accumulation of both monomeric and higher molecular weight (HMW) αSyn species in the Triton-insoluble fractions, in a manner depended on the endogenous αSyn and p25α protein load and fibril amount (Fig.S1 and S2). The recruitment of the endogenous oligodendroglial αSyn into pathological assemblies following treatment with the patient-derived fibrils was further confirmed by a thorough immunofluorescence analysis utilizing the rodent-specific D37A6 antibody to visualize the seeding of the endogenous rodent oligodendroglial αSyn, the human LB-509 to detect the exogenously added human material, the SYN303 to recognize the oxidized/nitrated αSyn, the MJFR-14-6-4-2 to monitor the aggregated αSyn and the EP1536Y antibody to observe the pathology-related phosphorylated at Serine 129 αSyn (pSer129-αSyn) (Fig.S3 and S4). Interestingly, human MSA-derived fibrils, unlike PD-derived ones, seem to exacerbate αSyn pathology in OLN-p25α cells, suggesting that a TPPP/p25α-enriched environment is more conducive to shaping different pathological αSyn conformations when disease-specific fibrils are introduced.

Having established the formation of intracellular pathological αSyn species within oligodendrocytes, following their inoculation with patient-derived fibrils, we aimed to decipher whether such species may be released extracellularly by the respective EVs and subsequently be implicated in the propagation of αSyn-related pathology. EM analysis verified the integrity and the morphological features of the oligodendroglial-derived EVs, which were enriched in concave particles enclosed by a phospholipid bilayer characteristic of exosomes and NTA confirmed the presence of particles with a size distribution pattern that corresponds to that of exosomes (Fig.1A-B). Although all fibril-treatments resulted in a comparable size distribution range between 40 to 170 nm, slight shifts in peak particle diameter between the different fibril-treatments and incubation times were observed. Moreover, the amount of EVs isolated from all fibril-inoculated (MSA1, PD, PFFs) OLN lines displayed a decrease at 48h post-fibril addition, compared to those released from control PBS-treated cells. Usually, this decrease was partially restored to the respective PBS-treated levels at 8 days post-treatment, indicating that EV release may be influenced by treatment duration (Fig.1B, MSA2-related data align with those in Fig. 1B). Moreover, under PBS conditions, OLN-p25α-derived EVs exhibited the highest concentration, with this difference being statistically significant when compared to OLN-ΑS7-derived ones. Equivalent trend of increased levels of OLN-p25α-derived EVs compared to those isolated from OLN-93 or OLN-AS7 cells was identified at 48h post-treatment with most fibril types (with an exception only in case of 6 μg/mL MSA1 or PD fibrils). Furthermore, the amount of OLN-93-derived EVs tended to be higher compared to OLN-AS7-derived upon treatment with 1 μg/mL MSA1, MSA2 fibrils and PFFs, whereas this relationship reversed when higher fibril concentrations were utilized. This observation differed though for PD fibrils. Finally, the results upon prolonged fibril incubation were not so clear, with EV concentrations varying depending on the fibril type and concentration employed (Fig.S5A).

**Figure 1:**
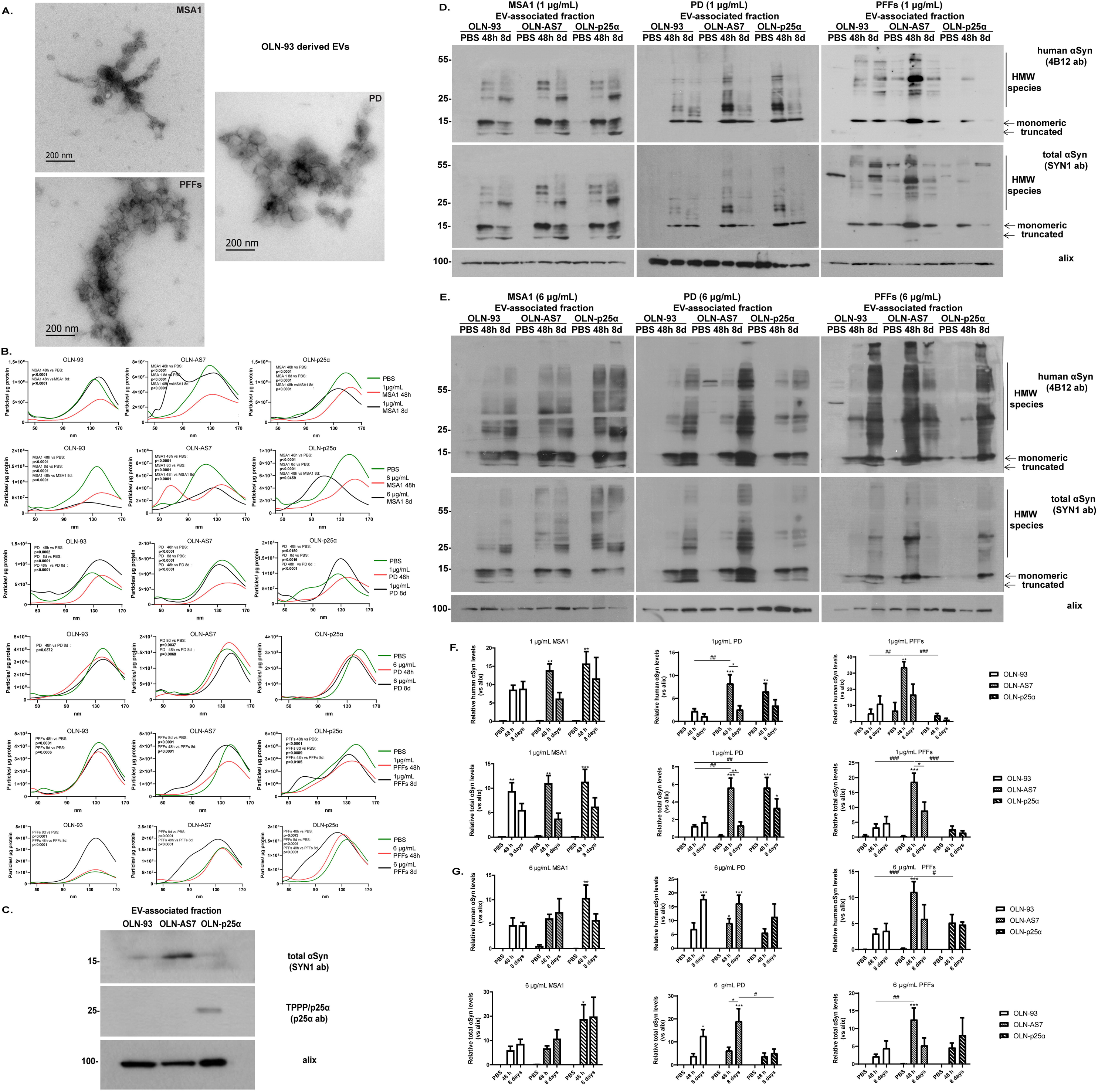
Human MSA or PD patient-amplified or recombinant αSyn fibrils alter the amount of oligodendroglial-secreted EVs and augment the release of EV-associated αSyn species. (A) Transmission EM images of OLN-93-derived EVs treated with MSA1, PD fibrils or PFFs (1 μg/mL, 48h). Scale bar: 200 nm. (B) Graphs depicting concentration and size distribution of EVs released from PBS-, MSA1-, PD- or PFF-treated oligodendrocytes (1 or 6 μg/mL, 48h or 8 days). (C) Immunoblots demonstrating human αSyn and TPPP/p25α protein presence within PBS-treated OLN-AS7- and OLN-p25α-derived ΕVs, respectively. (D-E) Immunoblots illustrating accelerated release of human and total αSyn via ΕVs derived from OLN cells treated with 1 (D) or 6 μg/mL (E) MSA1, PD fibrils or PFFs for 48h or 8 days. (F-G) Quantifications of human and total αSyn in EV-associated fractions of OLN cells treated with 1 (F) or 6 μg/mL (G) MSA1, PD fibrils or PFFs for 48h or 8 days. Alix confirms equal loading. Data are expressed as the mean ± SE of at least three independent experiments; *p< 0.05; **p < 0.01; ***p<0.001, by one-way ANOVA with Tukey’s post-hoc-test and #p<0.05; ##p<0.01; ### p<0.001, by two-way ANOVA with Bonferroni’s correction.

Under baseline conditions (PBS-treatment), both αSyn and TPPP/p25α proteins were released via EVs isolated from OLN-AS7 and OLN-p25α cells, respectively (Fig.1C). Importantly, in all OLN cell lines, the levels of EV-associated human and total αSyn were dramatically increased, following administration of all fibril types (MSA1, MSA2, PD, PFFs), in both fibril concentrations and time points assessed (Fig.1D-E, Fig.S5B-C). The secreted nanovesicles were specifically enriched not only in monomeric but also HMW and truncated αSyn species. Interestingly, their pattern following short-term incubation differed from the corresponding pattern of prolonged incubation and also varied among the different fibril types. Quantitative analysis indicated that the levels of EV-associated human and total αSyn reached a plateau at 48h upon addition of 1 μg/mL of all fibril types and then either remained unchanged or decreased over time in all OLN-cells (Fig.1F), with the exception of MSA2-treated OLN-cells (1 μg/mL), in which the levels of human and total αSyn not only remained high at both time points, but also increased over time particularly in OLN-p25α cells (Fig.S5D) Contrariwise, in the presence of higher amounts of MSA1 or PD fibrils and PFFs, αSyn levels either persisted or even increased at 8 days (Fig.1G). On the other hand, when 6 μg/mL of MSA2 fibrils were added, αSyn levels peaked at 48h and then declined (Fig.S5E). Interestingly, OLN-p25α-derived EVs contained statistically higher αSyn cargo, compared to OLN-93-derived at 48h post-MSA2 inoculation, whereas upon PD fibril- or PFF-treatment, OLN-AS7-derived EVs exhibited the highest αSyn levels. These results coincide with those obtained from the analysis of intracellular αSyn levels (Fig.S1-S4), further highlighting the importance of oligodendroglial αSyn and p25α protein load in the seeding of extracellular αSyn species and the heterogeneity in the response evoked by the different fibril types.To ascertain that the observed EV-associated αSyn species detected following fibril-treatment represent also protein species generated within oligodendrocytes and not only the human material that became associated with the outer EV membrane and subsequently co-precipitated during EV isolation, separate trypsin and saponine treatments were performed. In particular, PD-treated EVs (or PBS-treated as control) were incubated with trypsin, to eliminate any residual contamination with fibrils loosely attached to EVs. Trypsinized EV samples demonstrated a similar protein pattern with that of non-trypsinized (Fig. S5F), implying that a portion of αSyn species are produced within PD fibril-treated OLN cells and then released via EVs. Likewise, ELISA analysis revealed that saponine-treated EVs still contained significantly higher αSyn levels at 48h post-fibril addition (Fig.S5G), further suggesting the presence of αSyn species within the EV lumen.

### Human MSA and PD patient-derived fibrils or recombinant αSyn PFFs recruit the endogenous oligodendroglial αSyn, induce the phosphorylation of the seeded protein at Ser129 and augment the release of both αSyn conformations via oligodendroglial-derived EVs

To further characterize αSyn-related pathology engendered within OLN cells following patient-derived fibril addition, rodent seeded oligodendroglial αSyn (expressed at minimal levels under basal conditions [17]) and pathology-related pSer129-αSyn intracellular and EV-associated protein levels were examined. Addition of high patient-derived and recombinant αSyn fibril concentrations evoked the accumulation of endogenous rat αSyn in all OLN cells at both time points examined and, strikingly, enhanced its subsequent release by EVs (Fig.2A-B, Fig.S6A-B). By contrast, incubation with low fibril concentration failed to generate detectable signal (non presented data). Importantly, rodent αSyn was observed intracellularly only within the Urea-soluble fraction, where it appeared to be “trapped” in the stacking gel, similarly to its detection in the EV-associated fraction, pinpointing the rather insoluble nature of the seeded αSyn species engendered (Fig.2A-B, Fig.S6A-B). Interestingly, MSA fibril-treated OLN-p25α cells exhibited significantly higher levels of both intracellular and EV-associated rodent αSyn, compared to OLN-93 cells, whereas the respective signal upon PD fibril administration was more pronounced in OLN-AS7 cells, compared to the other cell lines (Fig.2C-D, Fig.S6C-D). Finally, in PFF-treated cells, the differences among the three lines did not lead to a safe conclusion. Immuno-EM analysis further verified the presence of both human (exogenously added) and rodent (endogenous seeded) αSyn within the lumen of OLN-p25α-derived EVs (Fig.2E). Regarding the temporal factors in the modulation of αSyn pathology, our results indicate that in general both intracellular and EV-associated rodent αSyn signal was detected at higher levels at 48h, compared to 8 days, post-fibril addition.

**Figure 2:**
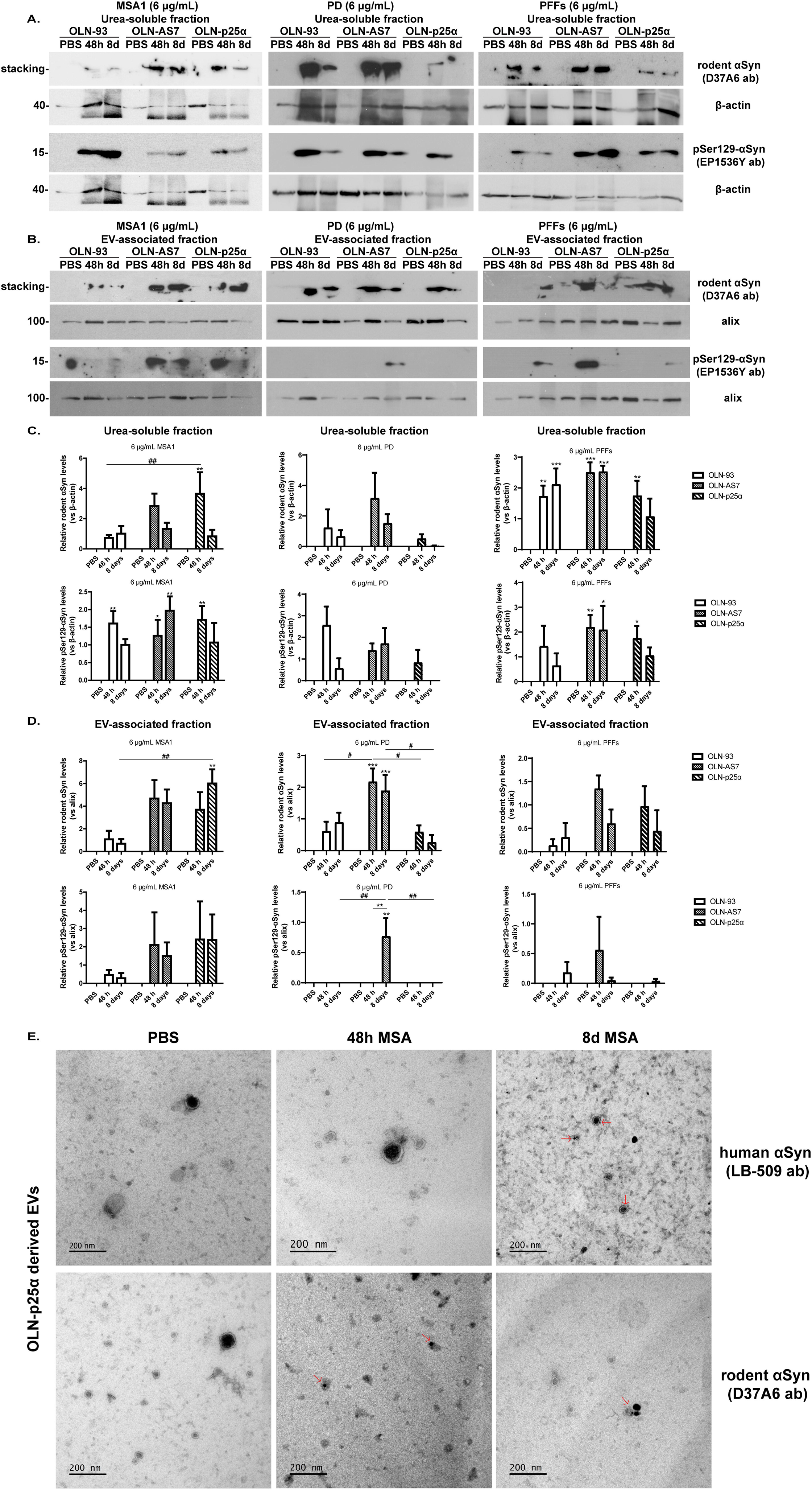
Incubation of rat oligodendroglial cells with high amounts of human MSA1, PD patient-derived fibrils or recombinant αSyn PFFs evokes the phosphorylation of the endogenous seeded αSyn at Ser129 and enhances its EV-associated release. (A-B) Representative immunoblots of rodent (endogenous) and pSer129-αSyn protein levels (D37A6 and EP1536Y antibodies, respectively) in the Urea-soluble (A) and EV-associated fractions (B) of OLN-93, OLN-AS7 and OLN-p25α cells treated with 6 μg/mL MSA1, PD fibrils or PFFs (or PBS as control) for 48h or 8 days. (C-D) Quantifications of the Urea-soluble (C) and EV-associated (D) rodent and pSer129-αSyn protein levels. Equal loading was verified by the detection of β-actin and alix. Data are expressed as the mean ± SE of at least three independent experiments; *p<0.05; **p<0.01; ***p<0.001, by one-way ANOVA with Tukey’s post-hoc-test and #p<0.05; ##p<0.01, by two-way ANOVA with Bonferroni’s correction. (E) Immuno-EM images of EVs, isolated from OLN-p25α cells treated with 6 μg/mL human MSA patient-amplified fibrils for 48h or 8 days (or PBS as negative control), indicate the presence of both human and rodent αSyn within the vesicle lumen (red arrows). Scale bar: 200 nm.

Similar to the rodent oligodendroglial αSyn, high amounts of intracellular pSer129-αSyn levels could be detected only in the Urea-soluble fraction at both time points and only following inoculation with high amounts of all fibril types (Fig.2A-B, Fig.S6A-B). No statistically significant association between pSer129-αSyn levels and different OLN cell type or time post-fibril addition was detected. Importantly, pSer129-αSyn produced within oligodendrocytes could be packed and released, at least partly, via EVs (Fig.2B&D), in a manner dependent on the fibril type harnessed. Specifically, in the presence of MSA1 fibrils, pSer129-αSyn was detected in all OLN cells, with OLN-AS7 and OLN-p25α cells expressing higher levels, compared to OLN-93, which sustained over time. In MSA2 fibril-treated cells, pSer129-αSyn signal was evident only at 48h and exclusively in OLN-AS7 and OLN-p25α cells. On the contrary, PD fibril-treatment was accompanied by the detection of EV-associated pSer129-αSyn only in OLN-AS7 and in particular at 8 days post-fibril addition. Finally, in PFF-treated OLN cells, EV-associated pSer129-αSyn was detected only in OLN-AS7-derived EVs at 48h and in all OLN cells at 8-days post-fibril addition (Fig.2C-D, Fig.S6C-D). The aforementioned data further underscore that the origin of αSyn fibrils—mainly oligodendroglial concerning MSA fibrils and neuronal in PD fibrils—may determine and differentiate, combined with the intracellular protein load, the manifestation of αSyn-related pathology within recipient cells.

### Addition of human MSA and PD patient-derived fibrils or recombinant αSyn PFFs to oligodendroglial cells stably overexpressing the human p25α protein influences both the levels and distribution of TPPP/p25α protein over time

We have previously shown that p25α is essential in modulating the formation of aberrant αSyn assemblies in OLN-p25α cells following treatment with recombinant αSyn PFFs [17]. Furthermore, the bidirectional relationship between αSyn and p25α burden suggests that changes in one may influence and facilitate the accumulation of the other [22, 23]. Consequently, understanding how p25α levels are regulated in response to different αSyn fibril strains could reveal potential strategies for modifying αSyn-related pathological outcomes in alpha-Synucleinopathies. Hence, changes in the protein levels and distribution (intracellularly and extracellularly) of TPPP/p25α protein, following incubation of OLN-p25α cells with MSA and PD patient-derived fibrils or PFFs (1 or 6 μg/mL for 48h or 8 days) were assessed. Consistently, treatment with all fibril types led to a reduction in the intracellular p25α levels (Triton-, SDS- and Urea-soluble fraction) at 48h post-fibril addition, compared to PBS-treatment (Fig.3A-F, left, Fig.S7A-B, left). However, the protein levels returned to baseline at 8 days. This reduction could be attributed either to enhanced protein secretion, in an attempt of the cell to discard excess p25α protein cargo, or induction of its degradation mechanisms as a response to proteolytic stress, or a combination of both scenarios. Consequently, we initially examined both EV-associated and non-EV-associated (EV-depleted fraction) extracellular p25α protein levels. Using both biochemical and immuno-EM analysis, we confirmed that the p25α protein is indeed present within EVs, in levels varying according to fibril type, concentration or treatment duration (Fig.3A-F, right, Fig3.I, Fig.S7A-B, right). Specifically, in MSA1-treated cells (1 μg/mL), a trend for decreased EV-associated p25α levels was observed at 48h, followed by a statistically significant increase at 8 days (Fig.3A), whereas in PD-treated cultures, EV-related p25α levels were significantly increased at 48h, with a similar trend at 8 days (Fig.3B), supporting the hypothesis of enhanced protein secretion. Conversely, in MSA2- (Fig.S7A) or PFF-treated (Fig. 3C) OLN-p25α cells, p25α levels within EVs decreased at 48h. Non-statistical significant differences were observed when higher fibril concentrations (6 μg/mL) were utilized (Fig. 3D-F).

**Figure 3:**
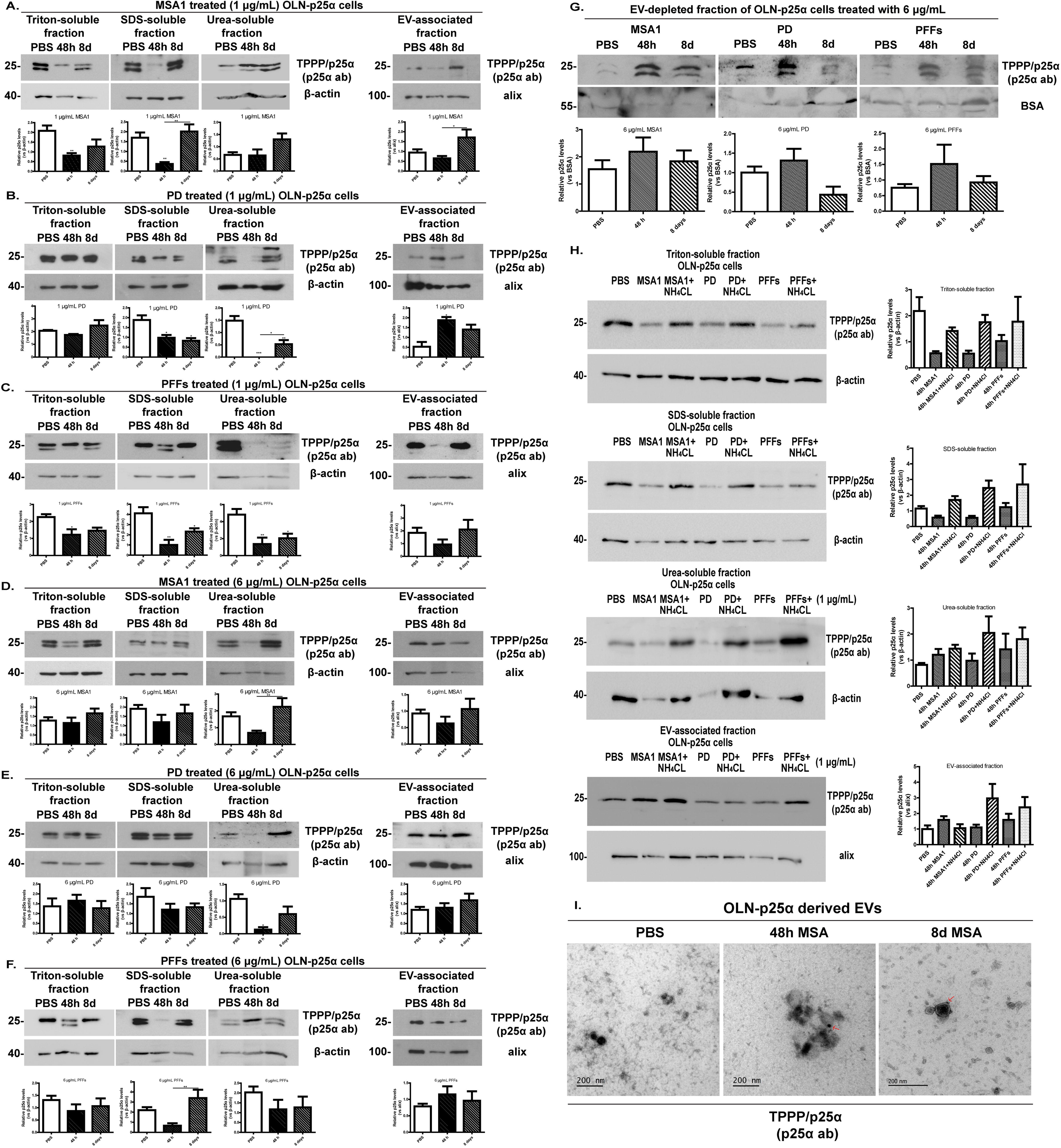
Addition of human MSA1 and PD patient-derived fibrils or recombinant αSyn PFFs alters the distribution of TPPP/p25α protein in a time-dependent manner. (A-F) Representative immunoblots and quantifications of Triton-, SDS-, Urea-soluble and EV-associated TPPP/p25α protein levels in OLN-p25α cells treated with 1 (Α,C,Ε) or 6 μg/mL (B,D,F) MSA1 (A-B), PD fibrils (C-D) or PFFs (E-F) (or PBS) for 48h or 8 days. β-actin and alix serve as loading controls. Data are expressed as the mean ± SE of three independent experiments; *p<0.05; **p<0.01; ***p<0.001, by one-way ANOVA with Tukey’s post-hoc-test. (G) Immunoblots of extracellular EV-free TPPP/p25α protein levels released from OLN-p25α cells treated with the different fibril types (6 μg/mL) or PBS for 48h or 8 days. Bovine Serum Albumin (BSA) confirms equal loading. Data are expressed as the mean ± SE of three independent experiments. (H) Immunoblots and quantifications of Triton-, SDS-, Urea-soluble and EV-associated TPPP/p25α protein levels from fibril-treated OLN-p25α cells (1 μg/mL) or PBS as control for 48h and then incubated with EV-depleted medium ± NH₄Cl. β-actin and alix ensure loading consistency. (I) Immuno-EM images of OLN-p25α-derived EVs treated with 6 μg/mL MSA1 fibrils for 48h or 8 days, verifying TPPP/p25α presence within EVs (red arrows). Scale bar: 200 nm.

Interestingly, non-EV-associated extracellular p25α levels were found increased in most fibril-treated OLN-p25α cells, 48h post-treatment (Fig.3G, Fig.S7C), suggesting accelerated excretion of p25α protein evoked by increased αSyn protein burden. To examine the second scenario, total lysosomal degradation was inhibited utilizing NH_4_Cl, based on our recent data suggesting that the lysosome is mostly responsible for p25α clearance in OLN-p25α cells [24]. Lysosomal inhibition reversed the changes on p25α levels in all fractions at 48h post-fibril addition, suggesting that αSyn fibrils indeed promote both the excretion and the lysosomal degradation of p25α (Fig.3H).

Finally, addition of both MSA and PD patient-amplified fibrils evoked a collapse in the p25α network and, more specifically, the well-shaped p25α protein pattern observed in PBS-treated cells acquired a fibrillar-like conformation, which also colocalized with the endogenous rodent and human αSyn, thus highlighting the incorporation of TPPP/p25α in pathological αSyn assemblies generated within oligodendrocytes following treatment with patient-derived αSyn fibrils (Fig.S7D).

### Exogenously added oligodendroglial EVs isolated from fibril-treated OLN cells are effectively taken up by primary rodent neuronal cultures and trigger the formation of αSyn-related pathological conformations, without disrupting neuronal network

Given that EVs from patient or recombinant αSyn fibril-treated oligodendrocytes are enriched in both αSyn and TPPP/p25α proteins, we investigated whether these disease-associated proteins can be taken up by primary rodent neuronal or oligodendroglial cultures. We initially labeled oligodendroglial-derived EVs with the PKH26 Red Fluorescent Cell Linker Mini Kit, which enabled the monitoring of EV uptake by recipient cells and subsequently applied the PKH26-labeled EVs to primary neuronal and oligodendroglial cultures. 3D-reconstruction of confocal microscopy images, created using Imaris software, illustrated that oligodendroglial-derived EVs were readily taken up by both neurons (TUJ1^+^, Fig.S8Α) and oligodendrocytes (p25α^+^, Fig.S8Α) following overnight incubation.

To evaluate the potential pathogenic propensity of oligodendroglial-derived EVs, primary mouse cortical cultures (from WT-αSyn P0-P2 pups) were exposed to EVs isolated from OLN-93, OLN-AS7 and OLN-p25α cells previously inoculated with 6 µg/mL human MSA1-, MSA2-, PD-fibrils or PFFs (or PBS as control) for 48h. Cultures were processed for immunocytochemical analysis 72 h or 6 days post-EV application, using antibodies against different αSyn species and/or pathological conformations, including the exogenously added human αSyn (reflecting the fibrils), along with the endogenous seeded rodent (of neuronal origin), oxidized/nitrated, aggregated and pSer129-αSyn.

Our results clearly suggest that EVs released from all fibril-treated OLN cells deliver human EV-associated αSyn in primary neurons (Fig.4A, Fig.S8B), with cultures inoculated with MSA1-treated OLN-p25α-derived EVs exhibiting the highest levels, and PD-treated OLN-p25α-derived EVs expressing the lowest levels (Fig.4A and 4E). Importantly, delivery of these EVs increased the levels of endogenous rodent neuronal αSyn (Fig.4B, Fig.S8C) and facilitated the generation of pathological αSyn assemblies (oxidized/nitrated, aggregated, pSer129-αSyn) within recipient cells (Fig.4C-D, Fig.S8D-F, Fig.5). Similar to human αSyn levels, neurons incubated with MSA1-treated OLN-p25α-derived EVs exhibited statistically significant higher oxidized/nitrated αSyn levels, compared to cultures incubated with fibril-treated OLN-93- or OLN-AS7-derived EVs (Fig.4E). Additionally, worth mentioning is that pSer129-αSyn could be detected only following prolonged incubation of primary cultures with these nanovesicles. Most importantly though, pSer129-αSyn^+^ signal was more potent when MSA1- and PFF-treated OLN-p25α-derived EVs were utilized, compared to the equivalent OLN-93-derived ones, further underpinning the contribution of TPPP/p25α protein in the establishment of MSA-like pathology (Fig.5D). In contrast, the presence of TPPP/p25α, when PD fibrils were utilized, did not accelerate αSyn-related pathology, as manifested by the lower pSer129-αSyn^+^ signal, as compared with MSA1- or PFF-treated EVs (Fig.5D).

**Figure 4:**
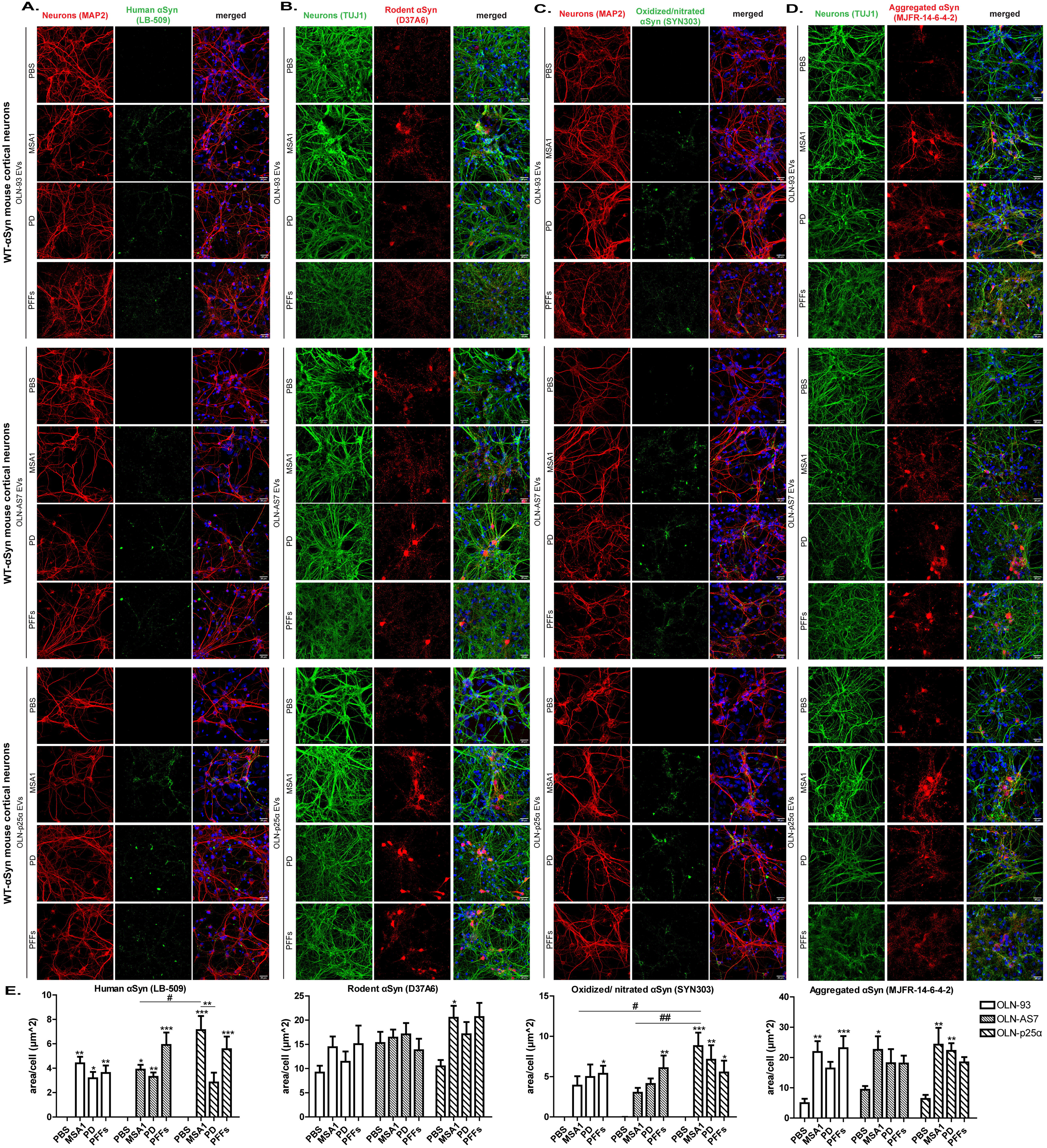
EVs released from OLN cells inoculated with human MSA- and PD-patient derived fibrils or recombinant PFFs carry aberrant αSyn conformations that are taken up by primary cortical neurons and trigger the aggregation of endogenous neuronal αSyn into pathological assemblies. Representative immunofluorescence images using antibodies against (A) neuronal dendrites (red, MAP2), human αSyn (green, LB-509), (B) neuronal axons (green, TUJ1), rodent αSyn (red, D37A6), (C) neuronal dendrites, oxidized/nitrated αSyn (green, SYN303), (D) neuronal axons, aggregated αSyn (red, MJFR-14-6-4-2) and DAPI staining in mouse primary cortical neurons inoculated with EVs (40 µg/mL for 72 h) derived from OLN cells previously treated with 6 µg/mL MSA1, PD fibrils or PFFs for 48h (or PBS as control). Scale bar: 25 µm. (E) Quantifications of human, rodent, oxidized/nitrated and aggregated αSyn protein levels in mouse primary neurons measured as area surface/cell following incubation with OLN-derived EVs. Data are expressed as the mean ± SE of three independent experiments with triplicate samples/condition within each experiment; *p<0.05; **p<0.01; ***p<0.001, by one-way ANOVA with Tukey’s post-hoc-test and #p<0.05; ##p<0.01, by two-way ANOVA with Bonferroni’s correction.

**Figure 5:**
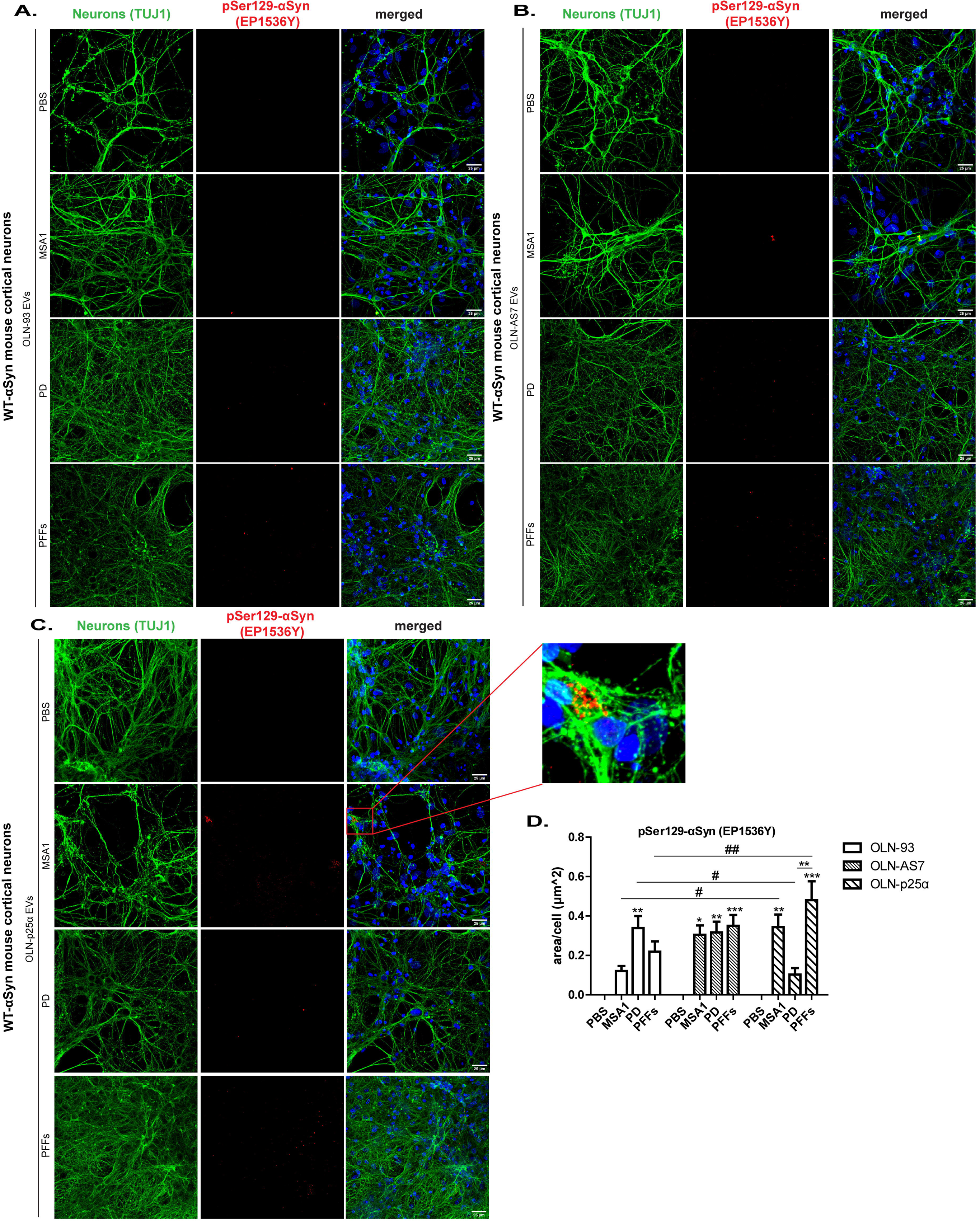
Oligodendroglial EVs released from human MSA1- and PD-patient amplified αSyn fibril- or PFF-treated OLN cells evoke the phosphorylation of αSyn at Ser129 within mouse primary cortical neurons. (A-C) Representative immunofluorescence images using antibodies against neuronal axons (green, TUJ1), pSer129-αSyn (red, EP1536Y) and DAPI staining in mouse primary cortical neurons incubated with EVs (40 µg/mL for 6 days) derived from OLN-93 (A), OLN-AS7 (B) and OLN-p25α (C) cells previously treated with 6 µg/mL MSA1- and PD-patient-derived fibrils or PFFs for 48h. Scale bar: 25 µm. (C) Zoomed image depicting pSer129-αSyn within TUJ1^+^ neurons following addition of EVs isolated from OLN-p25α cells treated with MSA1-patient-derived fibrils. (D) Quantification of pSer129-αSyn protein levels in mouse primary cortical cultures measured as area surface/cell following incubation with OLN cell-derived EVs. Data are expressed as the mean ± SE of three independent experiments with triplicate samples/condition within each experiment; *p<0.05; **p<0.01; ***p<0.001, by one-way ANOVA with Tukey’s post-hoc-test and #p<0.05; ##p<0.01, by two-way ANOVA with Bonferroni’s correction.

Collectively, such data indicate that oligodendroglial-derived EVs act as «Trojan horses» through which αSyn-related pathology transports in recipient cells, thus facilitating the propagation of disease-related species. To clarify whether αSyn-loaded EVs are sufficient to induce pathology in an αSyn null background, or the presence of endogenous rodent αSyn is required for the development of the disease-associated alterations, providing thus a clearer understanding of how the pathological cascade may be initiated, cortical neuronal cultures derived from KO-αSyn mice lacking endogenous αSyn expression were utilized. As illustrated in Fig. S9A-C, EV-associated αSyn can induce pathology alone, as both human and oxidized/nitrated αSyn species were detected in EV-treated KO-αSyn primary neuronal cultures. However, the levels of the formed αSyn species were lower, underpinning that while EVs trigger some level of pathology on their own, the presence of endogenous αSyn likely enhances or accelerates the aggregation and spread of pathological αSyn. This may suggest a synergistic relationship between EV-delivered and endogenous αSyn in driving MSA-like pathology.

Quantitative analysis of TUJ1^+^ (neuronal axon marker) and MAP2^+^ (neuronal dendrite marker) signals, expressed as the area occupied by each signal normalized to the cell number did not reveal significant impairment of either neuronal markers (Fig.S9D-G), implying that the neuronal network integrity was maintained, at least under the experimental conditions examined.

### Administration of fibril-treated oligodendroglial-derived EVs to rat primary oligodendroglial cultures recruit the endogenous oligodendroglial αSyn into the formation of pathological αSyn species, without altering the MBP^+^ network

Building on the internalization of PKH26-labeled oligodendroglial-derived EVs by primary rat oligodendroglial cultures (Fig.S10A) and the aforementioned neuronal αSyn-related pathology induced by fibril-treated oligodendroglial-derived EVs (Fig.4-5 and Fig.S8), we adopted a similar experimental approach to explore the potential pathogenic effects of these vesicles in primary rat oligodendroglial cultures.

As shown in Fig. 6A and S10B, oligodendroglial-derived EVs isolated from patient-derived (MSA1, MSA2, PD) or PFF-treated OLN cells, when introduced to primary rat oligodendroglial cultures, evoked the recruitment of rodent oligodendroglial αSyn, typically undetectable under normal conditions (PBS-EVs). Quantification analysis uncovered that regarding MSA fibrils, MSA1-treated OLN-p25α and MSA2-treated OLN-AS7 EVs exhibited the highest rodent αSyn levels when compared to MSA-treated OLN-93-derived EVs (Fig.6D, Fig.S10E). Furthermore, oxidized/nitrated αSyn species were formed (Fig.6A), but most importantly their signal co-localized with the endogenous oligodendroglial αSyn, underscoring its integration into pathological assemblies. Protein levels of oxidized/nitrated αSyn were significantly higher in PD- or PFF-treated OLN-AS7-derived EVs than OLN-93-derived (Fig.6D). Exogenous human αSyn was also detected (Fig.6B, Fig.S10C-E), with elevated levels following PFF EV-treatment isolated from all OLN cell lines. Aggregated αSyn conformations were also observed (Fig.6B, Fig.S10C), with significant differences noted only for MSA2 fibrils, where OLN-AS7 EVs manifested the highest levels (Fig.S10C-E). Importantly, fibril-treated EVs induced pathological phosphorylation of αSyn at Ser129 in primary rat oligodendrocytes (Fig.6C, Fig.S10D-E), with the strongest signal observed following addition of PFF-treated EVs isolated from all OLN lines. Statistically significant differences were detected when comparing MSA1-, PD-, and PFF-treated OLN-p25α EVs (Fig.6D). Additionally, MSA2-treated OLN-p25α EVs produced higher pSer129-αSyn levels than OLN-93 (Fig.S10E).

**Figure 6:**
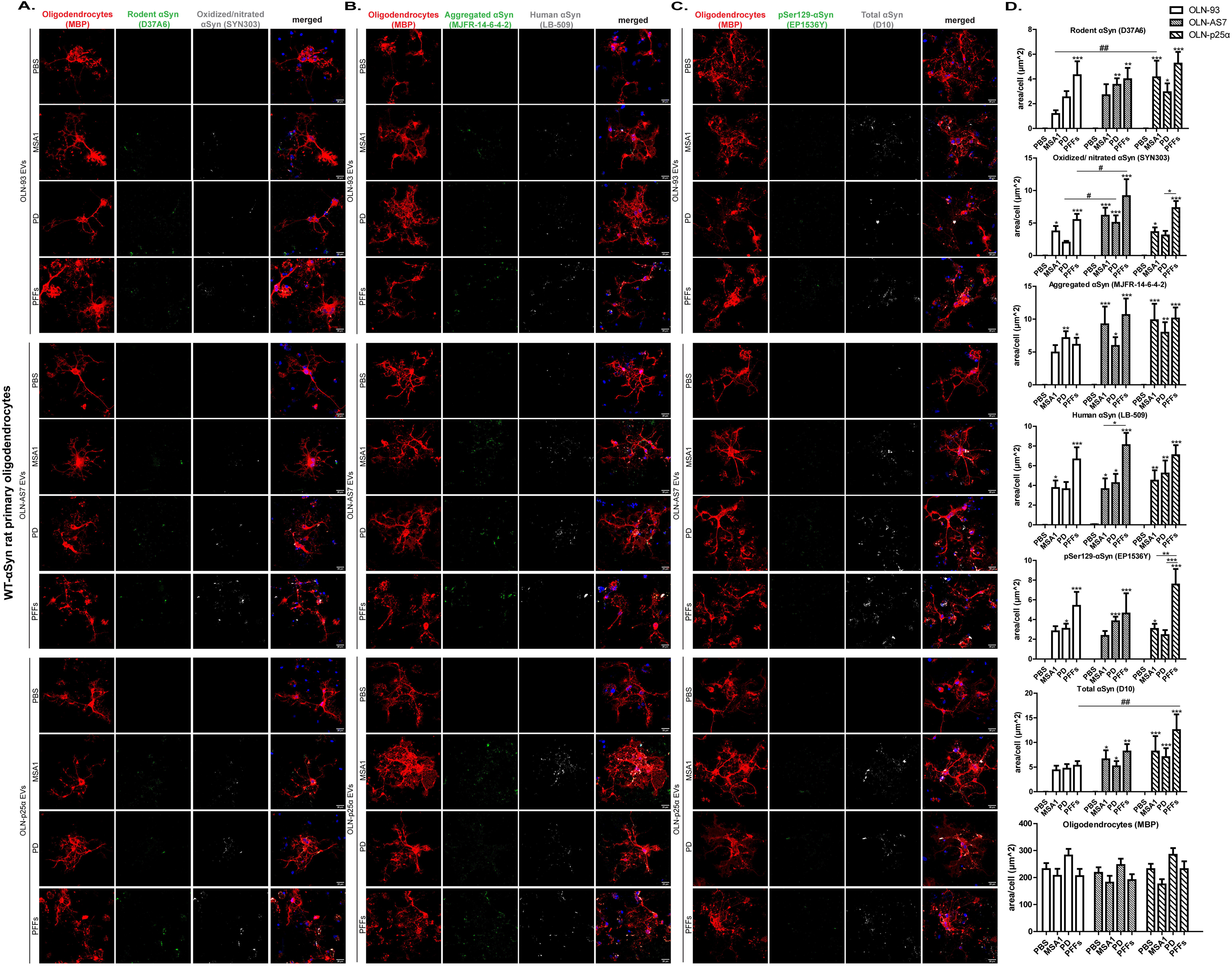
Oligodendroglial EVs released from human MSA1- and PD-patient derived fibril- or PFF-treated OLN cells recruit endogenous oligodendroglial αSyn and lead to the generation of pathology-related αSyn conformations within rat primary oligodendroglial cultures, without concomitant MBP-network disruption. (A-C) Representative immunofluorescence images using antibodies against (A) oligodendrocytes (red, MBP), rodent αSyn (green, D37A6), oxidized/nitrated αSyn (grey, SYN303), (B) oligodendrocytes, aggregated αSyn (green, MJFR-14-6-4-2), human αSyn (grey, LB-509), (C) oligodendrocytes, pSer129-αSyn (green, EP1536Y), total αSyn (grey, D10) and DAPI staining in rat primary oligodendrocytes inoculated with EVs (40 µg/mL for 72 h or 6 days for pSer129-αSyn) derived from OLN cells previously treated with 6 µg/mL MSA1- or PD patient-derived fibrils or PFFs for 48h (or PBS as control). Scale bar: 25 µm. (D) Quantifications of rodent, oxidized/nitrated, aggregated, human, pSer129 and total αSyn, as well as MBP protein levels in rat primary oligodendroglial cultures measured as area surface/cell following incubation with OLN cell-derived EVs (fibril- or PBS-treated). Data are expressed as the mean ± SE of at least three independent experiments with triplicate samples/condition within each experiment; *p<0.05; **p<0.01; ***p<0.001, by one-way ANOVA with Tukey’s post-hoc-test and #p<0.05; ##p<0.01, by two-way ANOVA with Bonferroni’s correction.

Finally, quantitative analysis of the area per cell covered by MBP (mature oligodendrocytes marker) did not reveal any statistically significant alterations between PBS- and fibril-treated conditions (Fig.6D, Fig.S10E). This highlights that the MBP^+^ network remained relatively unaffected despite the presence of pathological αSyn forms within myelinating oligodendrocytes, at least under the time frame assessed.

### EVs isolated from murine brains contain both αSyn (including the pSer129 pathology-related conformation that is highly enriched in PLP-hαSyn-derived EVs) and TPPP/p25α proteins and are efficiently delivered within the mouse dopaminergic system following unilateral intrastriatal injections

Following the thorough characterization of oligodendroglial-derived EVs and their seeding capacity in primary rodent cultures, we proceeded to the analysis of EVs isolated from different αSyn mouse models, including αSyn knockout mice (KO-αSyn), wild-type mice expressing endogenous αSyn (WT-αSyn) and transgenic mice expressing human αSyn selectively in oligodendrocytes (PLP-hαSyn), a well established MSA animal model. We aimed to characterize the EV-associated cargo in the three different genotypes and, subsequently, to evaluate the pathogenic capacity of PLP-hαSyn-originating EVs, in particular, *in vivo*. pSer129-αSyn DAB-immunostaining in brain regions encompassing the olfactory bulb, prefrontal cortex, striatum, hippocampus, substantia nigra pars compacta (SNpc) and cerebellum derived from six month-old WT-αSyn and PLP-hαSyn mice, revealed widespread pSer129-αSyn expression in the PLP-hαSyn mouse brain, predominantly in oligodendrocyte-abundant areas (Fig.S11), reinforcing the relevance of this model for studying MSA-related pathophysiological mechanisms. Afterwards, we isolated brain-derived EVs from the aforementioned mouse models, utilizing a sucrose gradient to enrich our EV preparations in exosomes, as described [19]. Notably, both αSyn and TPPP/p25α proteins, the two key GCI components accumulating in human MSA brains, were identified within these EVs (Fig.7A-B). Importantly, human and pSer129-αSyn were exclusively enriched in EVs derived from PLP-hαSyn brains, while endogenous rodent and total αSyn were present in EVs from both WT-αSyn and PLP-hαSyn brains, but absent in KO-αSyn-derived EVs. TPPP/p25α was detected at comparable levels across all mouse brain-derived EVs (Fig.7A-B).

**Figure 7:**
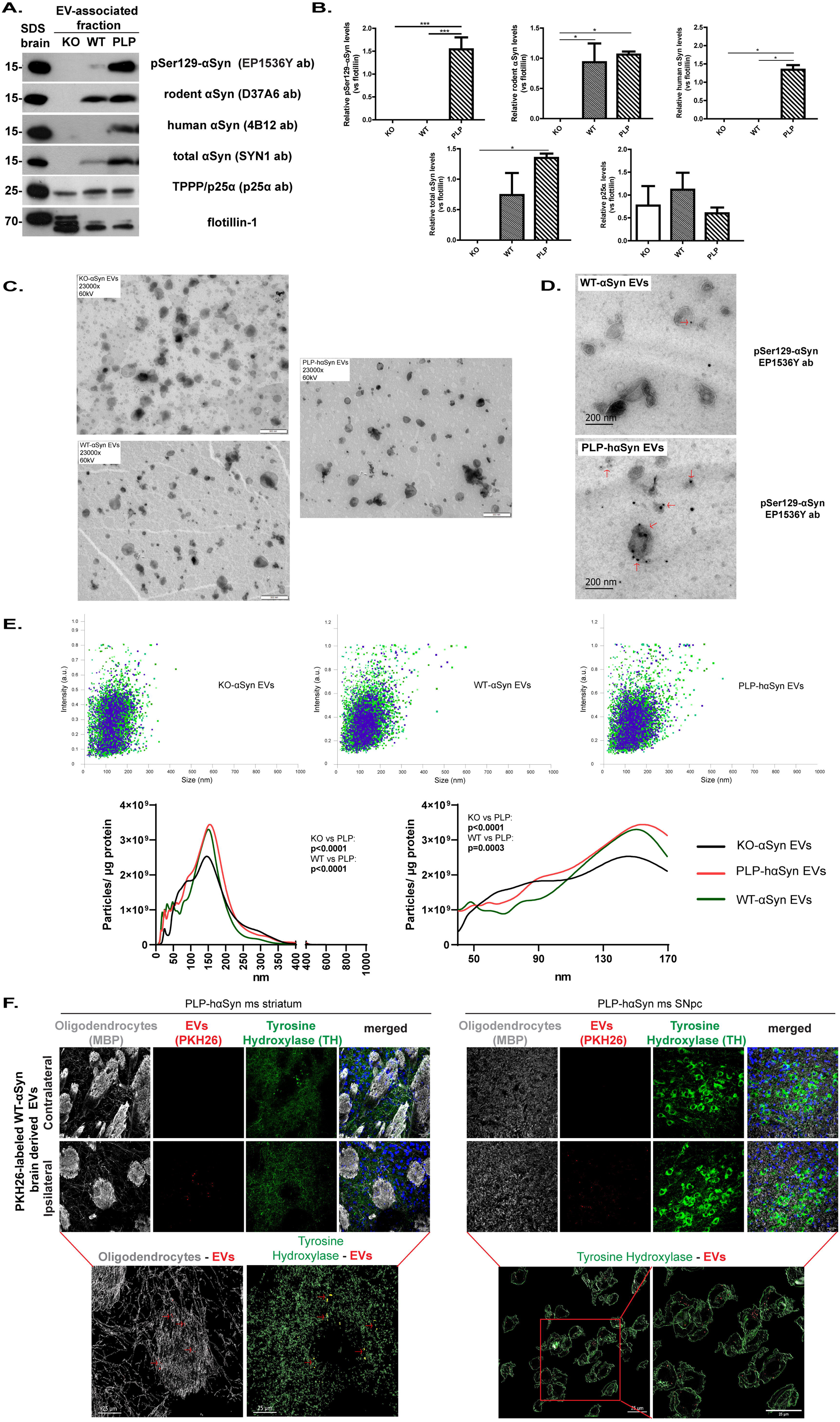
Characterization and efficient delivery of murine brain-derived EVs within the mouse dopaminergic system. (A) Representative immunoblots of pSer129 (EP1536Y), rodent (D37A6), human (4B12) and total αSyn (SYN1), and TPPP/p25α protein levels in the EV-associated fractions of six-month-old KO-αSyn, WT-αSyn and PLP-hαSyn mouse brains. (B) Quantifications of EV-associated pSer129, rodent, human and total αSyn, and ΤPPP/p25α proteins isolated from KO-αSyn, WT-αSyn and PLP-hαSyn mouse brains. Flotillin-1 served as EV loading control. Data are expressed as the mean ± SE of three independent experiments (n=3 mice/genotype); *p<0.05; **p <0.01; ***p<0.001, by one-way ANOVA with Tukey’s post-hoc-test. (C) Transmission EM images of EVs from KO-αSyn, WT-αSyn and PLP-hαSyn mouse brains (red arrows). Scale bar: 500 nm. (D) Immuno-EM images depicting increased pSer129-αSyn^+^ signal within PLP-hαSyn brain-derived EVs. Scale bar: 200 nm. (E) NTA confirmed the enrichment of brain-derived EVs in nanovesicles within the range of exosomes and uncovered higher particle concentration in PLP-hαSyn brain-derived EVs. (F) Efficient delivery of PKH26-labeled WT-αSyn brain-derived EVs in the mouse dopaminergic system detected at 7 days following unilateral intrastriatal injections. Immunofluorescence images depicting EV uptake by oligodendrocytes (MBP) and tyrosine-hydroxylase-positive (TH) terminals and neurons in the ipsilateral striatum and SNpc, respectively, further verified via 3D-reconstructed confocal imaging.

Transmission EM images verified the morphological integrity of the brain-derived EVs isolated from all three genotypes, with no apparent structural differences (Fig.7C) and immuno-EM analysis confirmed the enrichment of PLP-hαSyn-derived EVs in the pathology-related pSer129-αSyn (Fig.7D), consistent with the immunoblot data (Fig.7A-B). NTA revealed a similar size distribution profile across all genotypes, with the majority of EVs falling within the 100-200 nm range characteristic of exosomes. Interestingly, KO-αSyn mice exhibited statistically significant lower brain-derived EV amounts, as compared to PLP-hαSyn that expressed the highest levels (Fig.7E). Such data may insinuate that endogenous αSyn protein load may influence EV biogenesis and/or release or that regulatory mechanisms affecting EV levels may vary among different genotypes.

To further investigate the pathogenic potential of the aforementioned nanovesicles *in vivo*, EVs isolated from WT-αSyn brains were labeled with the fluorescent dye PKH26 and subsequently injected into the right dorsal striatum of PLP-hαSyn mice. Confocal microscopy analysis clearly demonstrated the presence of PKH26-labeled EVs within the ipsilateral striatum, seven days post-EV injection (Fig.7F). This signal co-localized with both MBP and TH, indicating nanovesicle uptake by both oligodendrocytes and dopaminergic axons (Fig.7F). Additionally, EV-positive signal was detected within TH^+^ dopaminergic neurons within the ipsilateral SNpc, suggesting EV transfer along the nigrostriatal axis, as additionally validated through 3D reconstruction (Fig.9F).

### Intrastriatal inoculation of PLP-hαSyn and not WT-αSyn brain-derived EVs triggers the formation of pSer129-αSyn^+^ conformations along the mouse nigrostriatal axis, without evident dopaminergic system degeneration

To investigate the potential contribution of αSyn-enriched EVs, preferentially of oligodendroglial origin, in the spread of MSA-like pathology, WT-αSyn mice were inoculated with EVs isolated from either WT-αSyn (negative control) or PLP-hαSyn (positive control) mouse brains. Our data demonstrate that PLP-hαSyn-derived EVs induced significant pSer129-αSyn^+^ pathology in the injected striatum, co-localizing with both mature oligodendrocytes (MBP^+^) and dopaminergic neuron terminals (TH^+^), whereas no pathology was detected in the corresponding hemisphere of mice inoculated with WT-αSyn EVs (Fig. 8A). The detected pSer129-αSyn^+^ signal increased over time, indicating a time-dependent pattern of pathology propagation (Fig. 8B). Furthermore, pSer129-αSyn-related pathology spread to the contralateral hemisphere at three-month post-EV injection, probably suggesting trans-synaptic transmission (Fig. 8A-B). Importantly, pSer129-αSyn-related pathology propagated to the SNpc, as manifested by the detection of pSer129-αSyn^+^ signal within TH^+^ dopaminergic neurons of the ipsilateral SNpc (Fig. 8C). This accumulation was more pronounced at three months post-injection, highlighting again a time-dependent progression of pathology (Fig. 8D).

**Figure 8:**
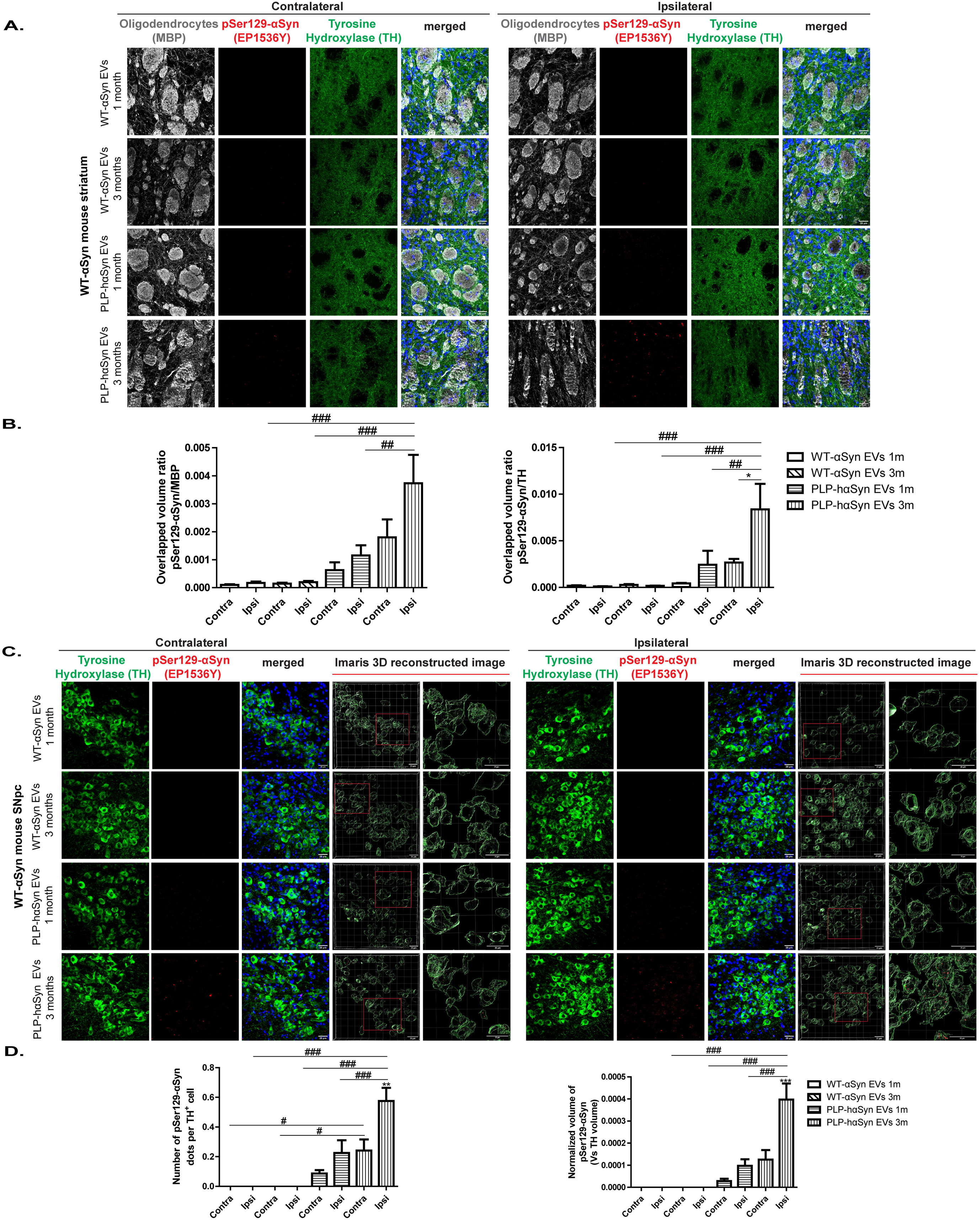
Inoculation of WT-αSyn mice with EVs isolated from PLP-hαSyn and not WT-αSyn mice, is accompanied by the formation of pSer129-αSyn^+^ inclusions that increase over time. (A) Representative immunofluorescence striatal images depicting mature oligodendrocytes (grey, MBP), pSer129-αSyn (red, EP1536Y), tyrosine-hydroxylase (green, TH) and DAPI in WT-αSyn mice injected with WT-αSyn or PLP-hαSyn EVs at one or three months. Scale bar: 25 µm. (B) Quantification of pSer129-αSyn^+^overlapped volume ratio with MBP^+^ or TH^+^ signal following mouse brain-derived EVs intrastriatal delivery. Data are expressed as the mean ± SE (n=6 animals/group); *p<0.05, by one-way ANOVA with Tukey’s post-hoc-test, comparing between hemispheres and ##p<0.01; ###p<0.001, by two-way ANOVA with Bonferroni’s correction, comparing between EV-groups. (C) Representative immunofluorescence ventral midbrain images of pSer129-αSyn, TH and DAPI in WT-αSyn mice injected with WT-αSyn or PLP-hαSyn EVs at one or three months. 3D-reconstruction confirms pSer129-αSyn presence within dopaminergic neurons. Scale bar: 25 µm. (D) Quantification of pSer129-αSyn dots per TH^+^ cell and pSer129-αSyn volume normalized to total TH volume following mouse brain-derived EVs delivery. Data are expressed as the mean ± SE (n=6 animals/group); **p<0.01; ***p<0.001, by one-way ANOVA with Tukey’s post-hoc-test, and #p<0.05; ###p<0.001, by two-way ANOVA with Bonferroni’s correction.

Despite the observed pSer129-αSyn pathology, no significant differences in dopaminergic terminal density or TH^+^ neuron cell counts between hemispheres (EV-injected or not) or EV groups (derived from WT-αSyn or PLP-hαSyn mouse brains) were observed (Fig.S12). This suggests that pSer129-αSyn accumulation does not immediately compromise dopaminergic system integrity, at least within the time frame examined.

### Oligodendroglial EVs originating from OLN-p25α cells treated with human MSA patient-amplified αSyn fibrils induce pSer129-αSyn accumulation along the nigrostriatal axis of recipient PLP-hαSyn mice

Considering the aforementioned data and in order to elaborate further the contribution of oligodendroglial-derived EVs to the development and progression of MSA-related pathology *in vivo*, we focused on the three-month timepoint, where αSyn-pathology was most pronounced and utilized PLP-hαSyn mice, instead of WT-αSyn, as hosts. As depicted in Fig. 9Α-Β, intrastriatal administration of EVs isolated from OLN-p25α cells pre-incubated with human fibrils derived from MSA patient brains led to pSer129-αSyn accumulation in the injected hemisphere and facilitated the spread of αSyn-associated pathology throughout the rostrocaudal axis encompassing the striatum. Moreover, co-localization of pSer129-αSyn^+^ dots with MBP^+^ and TH^+^ signal, was significantly augmented, as compared to the control EV PBS-treated group (Fig. 9C). However, the volume occupied by MBP and TH signals did not differ among the four groups, indicating that both cell populations remained relatively unaffected despite the ongoing pathology (Fig. 9C).

**Figure 9:**
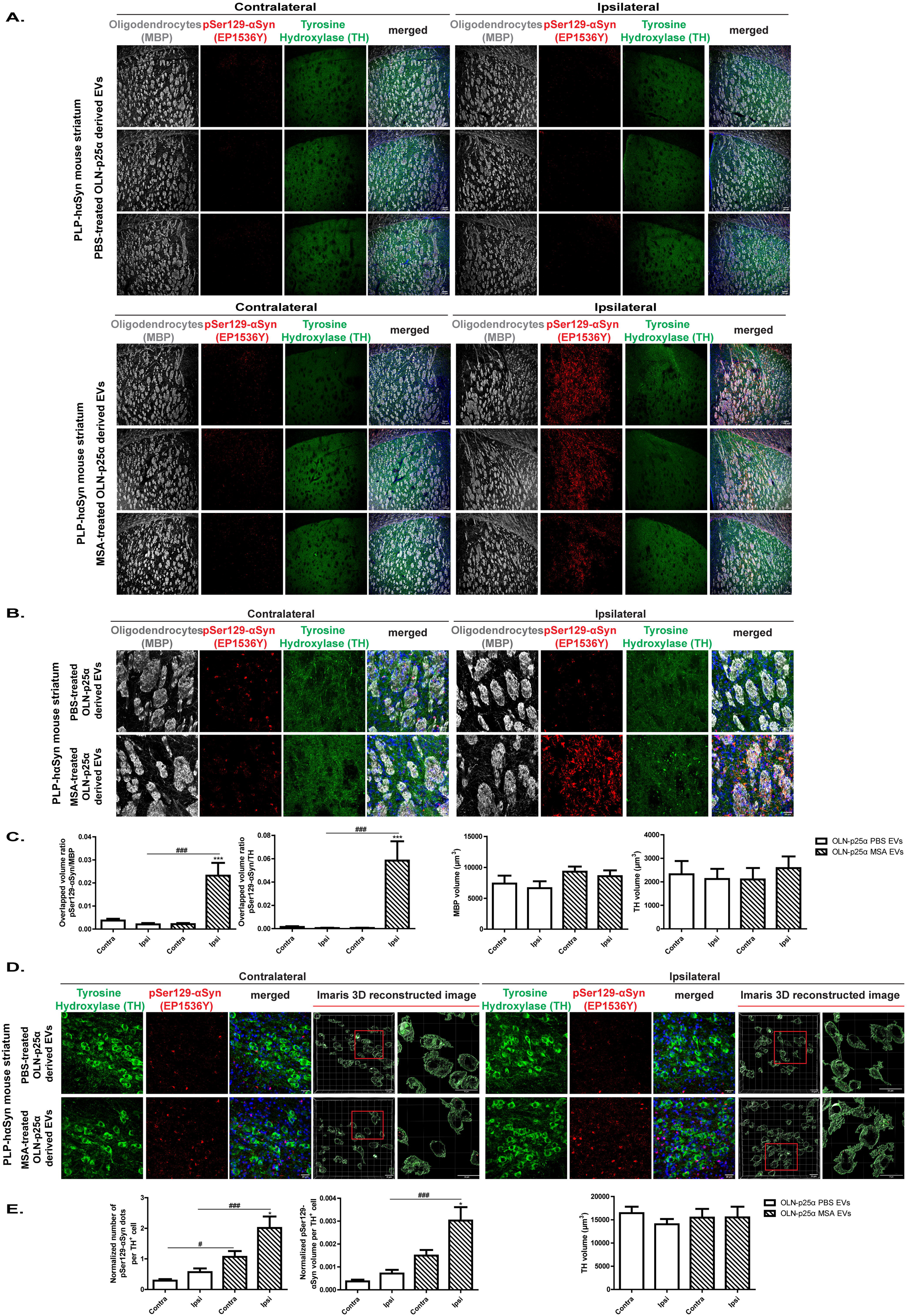
EVs isolated from oligodendroglial cells inoculated with human MSA patient-derived fibrils induce widespread pSer129-αSyn accumulation along the nigrostriatal axis of recipient PLP-hαSyn mice. (A-B) Representative immunofluorescence images of MBP^+^ oligodendrocytes (grey), pSer129-αSyn (red), TH^+^ neurons (green) and DAPI in PLP-hαSyn mouse striatum inoculated with PBS- or MSA-treated OLN-p25α EVs, three months post-injection. Scale bar: 100 µm (A) and 25 µm (B), respectively. (C) Quantifications of pSer129-αSyn^+^/TH^+^, pSer129-αSyn^+^/MBP^+^overlapped volume ratio, MBP^+^ or TH^+^ volume of PLP-hαSyn mouse striata inoculated with OLN-p25α-derived EVs. Data are expressed as the mean ± SE (n=3 mice/group); ***p<0.001, by one-way ANOVA with Tukey’s post-hoc-test and ###p<0.001, by two-way ANOVA with Bonferroni’s correction. (D) Representative immunofluorescence and 3D-reconstructed images of TH, pSer129-αSyn and DAPI in PLP-hαSyn mouse SNpc, three months following intrastriatal injections with PBS- or MSA-treated OLN-p25α EVs. Scale bar: 25 µm. (E) Quantifications of pSer129-αSyn^+^/TH^+^ cell, pSer129-αSyn+/total TH^+^ volume and total TH^+^ volume in the ventral midbrain of PLP-hαSyn mice inoculated with OLN-p25α-derived EVs. Data are expressed as the mean ± SE (n=3 mice/group); *p<0.05, by one-way ANOVA with Tukey’s post-hoc-test, and #p<0.05; ###p<0.001, by two-way ANOVA with Bonferroni’s correction.

Importantly, αSyn-related pathology propagated also to the ipsilateral SNpc of PLP-hαSyn mice injected with MSA-treated OLN-p25α EVs, thus highlighting the capacity of these EVs to template the endogenous αSyn into the formation of pathological protein conformations along the nigrostriatal axis (Fig. 9D). Both the number and volume of pSer129-αSyn^+^ signals within SNpc dopaminergic neurons were significantly higher in the hemisphere injected with the MSA-treated EVs, compared to the control PBS-treated one. Interestingly, a subtle increase of pSer129-αSyn^+^ signal was also observed in the non-injected SNpc, supporting the idea of trans-synaptic propagation (Fig. 9E).

Finally, dopaminergic system integrity remained relatively unaffected, as evidenced by the preserved striatal terminal density (Fig.S13Α-Β) and the TH^+^ nigral cell volume (Fig.S13C-D). Similarly, behavioral assessments (challenging beam test, rotarod) did not uncover significant motor impairments across examined groups (Fig.S13E).

### EVs isolated from human MSA brains augment nigrostriatal pSer129-αSyn pathology without compromising dopaminergic integrity or motor function of host PLP-hαSyn mice

To delve deeper into the contribution of EV-associated pathological αSyn to the spread and progression of MSA-like pathology *in vivo*, we isolated EVs from the brains of MSA patients or respective controls. ΝΤΑ revealed that EVs purified from MSA brain fractions (b, c, d) using a sucrose gradient displayed a distinct size distribution, with a predominant population within the 100-200 nm range, consistent with the typical size of exosomes (Fig.10A). Immunoblot analysis demonstrated that EVs from human MSA brain samples contained elevated αSyn levels, in particular HMW and truncated conformations, compared to control (CTR)-derived EVs (Fig.10B), suggesting enhanced release of pathological αSyn species via EVs in human MSA brains. Notably, for the first time TPPP/p25α protein was also detected in EV-associated fractions of both human MSA and CTR samples, with no significant differences among the groups (Fig.10Β). Although pSer129-αSyn was undetectable via immunoblot, immuno-EM analysis uncovered an increased pSer129-αSyn, enrichment of MSA-derived EVs, compared to CTR-derived EVs, present both in the membrane and the lumen of the vesicles, (Fig.10C).

**Figure 10:**
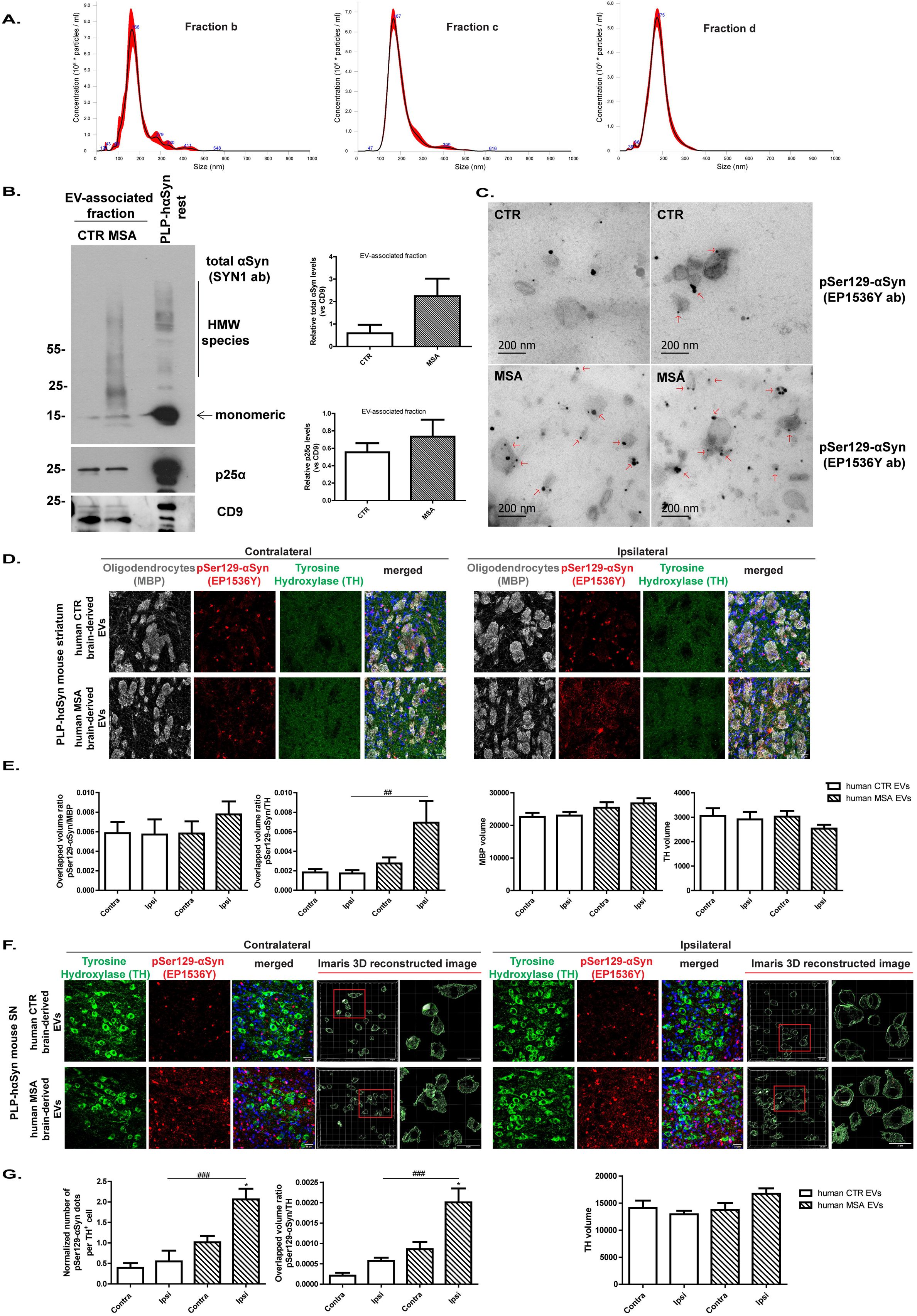
Brain-derived EVs isolated from MSA patients exacerbate pSer129-αSyn accumulation along the nigrostriatal axis of PLP-hαSyn mice. (A) NTA-generated size distribution plot of b, c and d human MSA brain-derived EV fractions isolated with a sucrose gradient. (B) Representative immunoblots and quantifications of αSyn (monomer, HMW, truncated) and TPPP/p25α proteins within MSA- or CTR-derived EVs. CD9 served as EV-loading control. (C) Immuno-EM images demonstrating pSer129-αSyn enrichment (red arrows) within MSA-derived EVs. Scale bar: 200 nm. (D) Representative immunofluorescence images of MBP^+^ oligodendrocytes (grey), pSer129-αSyn (red), TH (green) and DAPI in the PLP-hαSyn EV-injected striatum, three-months post-injection. Scale bar: 25 µm. (E) Quantifications of the overlapped volume ratio of pSer129-αSyn^+^/MBP^+^, pSer129-αSyn^+^/TH^+^, MBP^+^ or TH^+^ volume in human EV-injected PLP-hαSyn striatum. (F) Representative immunofluorescence and 3D-reconstructed images of pSer129-αSyn, TH and DAPI in PLP-hαSyn nigra, three-months following human-EV intrastriatal injections. Scale bar: 25 µm. (G) Quantifications of nigral pSer129-αSyn^+^/TH^+^ cell, pSer129-αSyn^+^/TH^+^volume and total TH^+^ volume. Data in E and G are expressed as the mean ± SE (n=4 mice/group); *p<0.05, by one-way ANOVA with Tukey’s post-hoc-test and #p<0.05; ##p<0.01; ###p<0.001, by two-way ANOVA with Bonferroni’s correction.

Subsequently, the aforementioned EVs (n=2 MSA or 2 CTR donors) were unilaterally injected in the striatum of three-month old PLP-hαSyn mice. Our data clearly demonstrate that MSA-EV-treated mice exhibited significant accumulation of pSer129-αSyn^+^ signal within the ipsilateral striatum three-months post-EV inoculation, compared to the contralateral one (Fig.10D-E). Although a trend towards increased co-localization of pSer129-αSyn^+^ with MBP^+^ signals was observed in MSA-EV-treated mice (Fig.10E), it did not yet reach statistical significance, likely due to the limited sample size. Contrariwise, pSer129-αSyn^+^ co-localization with TH^+^ dopaminergic neurons was significantly higher in the ipsilateral MSA-EV-treated hemisphere, compared to the CTR-EV-treated one (Fig.10E). Importantly, no significant differences in MBP^+^ or TH^+^ volume were observed among groups, indicating preserved cell populations, at least at the timeframe examined (Fig.10E).

Similar to MSA fibril-treated OLN-p25α-derived EVs, *in vivo* administration of human MSA brain-derived EVs induced pSer129-αSyn-related pathology in the SNpc of PLP-hαSyn mice, demonstrating spread of pathology along the nigrostriatal axis, distal to the injection site (Fig.10F). Both the number and volume of pSer129-αSyn^+^ inclusions within the nigral dopaminergic neurons were significantly higher in MSA-EV-treated mice, compared to CTR-EV-treated ones (Fig.10G).

Finally, assessment of the dopaminergic system integrity by measuring striatal dopaminergic fiber density and nigral TH^+^ neuron numbers did not reveal significant differences between the contralateral and ipsilateral sides of both MSA patient and control EV-treated groups (Fig.S14A-D). Motor performance, assessed via the challenging beam and rotarod tests, was similar between groups (Fig.S14E), though additional samples are needed to draw safe conclusions.

## Discussion

This study underscores the pivotal role of EVs isolated from cell and animal MSA experimental models, as well as, from human MSA brains in the dissemination of αSyn-related pathology, while also highlighting the contribution of distinct fibril types (MSA versus PD or PFFs) in influencing pathology progression and characteristics. Levels of EV-associated αSyn and TPPP/p25α, the two main GCI components, were significantly altered following exposure to pathological αSyn fibrils amplified from human MSA or PD patient brains (or αSyn PFFs), in a manner dependent on the originating source (oligodendroglial versus neuronal αSyn fibril strains). Importantly, αSyn- and TPPP/p25α-enriched EVs were readily taken up by neurons and oligodendrocytes, driving αSyn propagation *in vitro* and *in vivo*. By compiling the data from the rodent MSA-like cellular and animal models with those obtained from human MSA brains, we aimed to link the cargo and properties of EVs released under experimental MSA-like and human MSA conditions with recipient cell responses and downstream effects on cellular and network integrity in the context of MSA.

Prevailing hypotheses underlying the aberrant intracellular aggregation of αSyn within MSA oligodendrocytes, suggest either excessive expression of the *SNCA* gene encoding for αSyn [25] or uptake of extracellular αSyn following its secretion by neighboring neurons [7]. During the last decades, a growing body of evidence pinpoints EVs, and exosomes in particular, as key players in αSyn intercellular transmission in PD, focusing mostly on neuronally-derived EVs [10, 12, 26–28]. However, the current literature possesses shortcomings regarding the contribution of oligodendroglial-derived EVs in the development and progression of MSA-related pathology, in conjunction with the lack of data regarding the role, if any, of extracellular EV-associated TPPP/p25α. Several studies have demonstrated that exosomes isolated from serum or plasma by immunoprecipitation assays using cell-specific markers carry cargo from their putative cells of origin through the blood–brain barrier and thus could provide a “*window*” into pathologic processes occurring in the brain, offering a major advancement in analyzing brain biomarkers non-invasively [28–31]. An initial study reported that neuronal-derived plasma EVs were statistically significantly lower in MSA compared to PD patients [32], whereas others suggested that αSyn cargo within neuronal-derived serum EVs in MSA subjects was unchanged compared to controls and concomitantly was two-fold less than that measured in PD patients [33, 34]. In regards to oligodendroglial-derived EVs, it has been proposed that both their concentration and associated αSyn cargo was lower in the MSA plasma as compared to healthy controls [14]. On the other hand, αSyn protein cargo was found elevated in putative oligodendroglial-derived EVs isolated from MSA blood plasma or serum compared to PD or controls, whereas neuronal-derived EVs exhibited higher αSyn levels in PD versus MSA plasma or serum [35], further supporting the oligodendroglial or the neuronal contribution to MSA and PD pathogenesis, respectively. Our NTA cell culture data suggest that administration of all fibril types (patient or recombinant) altered the EV amount, in a concentration- and time-dependent manner. Interestingly, PLP-hαSyn mice exhibited the highest brain-derived EV levels, as compared to those of WT-αSyn or KO-αSyn mice. How the ratio of αSyn protein cargo associated with oligodendroglial or neuronal EVs fluctuates in human MSA or PD brains remains to be determined.

Previous studies pointed out that oligodendroglial-derived exosomes can be taken up by neurons and exert multifaceted effects on neuronal function [36]. Under baseline conditions, the exosome-mediated communication between neurons and oligodendrocytes appears in terms of coordination of the myelin sheath formation [37, 38], or in terms of metabolic support of neurons and thus neuroprotection against oxidative stress and starvation [39]. The novelty of our approach lies in the elucidation of EV-mediated αSyn propagation driving oligodendroglial αSyn overload that may ultimately lead to oligodendroglial dysfunction and neuronal demise observed in MSA. The co-localization of pathological αSyn with TPPP/p25α within rodent- and brain-derived EVs further supports the hypothesis that oligodendroglial-derived vesicles operate as carriers of noxious protein conformations, templating the spread of pathological αSyn and/or TPPP/p25α species within recipient neuronal or oligodendroglial cells. Both the endogenous rodent oligodendroglial and the pathology-related pSer129-αSyn were enriched in oligodendroglial-derived EVs following incubation with MSA or PD patient fibrils (or PFFs as control). Brain-derived EVs from the PLP-hαSyn MSA mouse model and from human MSA brains, contained significant amounts of pSer129-αSyn, compared to respective controls, indicating a potential role of this conformation to the spread of pathology. Interestingly, and in support to the “strain hypothesis”, MSA fibrils triggered Ser129 αSyn phosphorylation selectively in OLN-p25α cells whereas PD fibrils only in OLN-AS7 cells, further highlighting the contribution of TPPP/p25α protein to MSA-like pathogenesis.

The observed propapagation of pathological αSyn conformations to both oligodendroglial and neuronal cells via oligodendroglial EVs aligns with studies on exosome-mediated αSyn spread in PD models [11, 40], suggesting a conserved role for EVs in αSyn transmission across alpha-Synucleinopathies. The validation of the seeding activity of these EVs demonstrated herein both in cell and animal models, highlights the importance of investigating oligodendroglial-derived EVs in particular as potential culprits for MSA pathogenesis. Such findings line up with prior studies that failed to detect increased αSyn protein load within L1CAM^+^ neuronal-derived EVs from MSA serum [33]. The fact that pathological (MJFR14^+^ oligomeric/filamentous) soluble αSyn conformers can be detected in blood plasma-derived neuronal EVs from PD patients and can be amplified with the seeding amplification assay [41] further underscores the potential use of peripheral EVs as disease biomarkers. Towards this direction, a recent study demonstrated that seeding activity of misfolded αSyn derived from neuronal exosomes in blood is inversely associated with PD diagnosis and disease duration, probably reflecting the neuronal loss and/or decline of proteolytic pathways in later disease stages [42].

However, several limitations must be acknowledged when interpreting the aforementioned results. This study relies on *in vitro* and murine models, which, while informative, do not fully recapitulate the complexity of the human disease. For example, differences in the EV composition and/or uptake mechanisms between rodents and humans may limit translational relevance. Furthermore, the dearth of longitudinal studies restricts our understanding of the cumulative impact of EV-induced pathology on neuronal survival and motor performance. While dopaminergic integrity was preserved within the three-month timeframe, prolonged αSyn accumulation could possibly lead to neurodegeneration, as observed in advanced disease stages. Finally, the narrow sample size, in experiments involving the human-derived EVs, requires further validation in larger cohorts to achieve reproducibility and statistical consistency.

To bridge these gaps and advance towards clinical translation, several steps are required. First, the development of humanized models incorporating patient-derived induced pluripotent stem cells or brain organoids would provide more accurate systems for evaluating EV-mediated pathology. Additionally, identifying EV biomarkers specific to MSA and PD could enable early diagnosis and patient stratification, similar to the strategies developed by [35, 43–48]. Moreover, understanding the differential contributions of neuronal and glial-derived EVs to disease mechanisms opens avenues for developing cell-type-specific interventions and patient stratification and personalized treatment in alpha-Synucleinopathies. Finally, therapeutic interventions targeting EV release [49], cargo loading [50, 51], or uptake should also be exploited in preclinical and clinical trials.

In conclusion, this study provides compelling evidence that oligodendroglial-derived EVs play a pivotal role in the progression of MSA, offering new avenues for understanding disease mechanisms and developing targeted interventions. By addressing the limitations outlined and leveraging emerging technologies, the field can move closer to translating these findings into effective treatments for MSA, PD, and related disorders.

## Acknowledgments

The authors would like to thank the Biological Imaging Facility at BRFAA, and especially Tasos Delis, for their valuable assistance with confocal imaging and image analysis. We also acknowledge the BRFAA Animal Facility for their support. Special thanks are extended to Katerina Melachroinou, Maria Fouka and Claudia Schwiegk for their exceptional technical assistance.

## Author contributions

MX conceived the idea of the study, secured the funding, and supervised the research. MX also contributed to the revision and final editing of the manuscript. MV was responsible for creating the visualizations and drafting the original version of the manuscript. MV, DD, PM, FA, GT, EML, MG, SF, AK, IT and SH contributed to the experimental work and data analysis, ensuring the reliability and accuracy of the results. VGG, NS, PHJ, MZ, and SB provided key resources, including the electron microscope facility, experimental animals, antibodies, cells and pathological fibrils, which were essential for the successful completion of the study.

## Data and materials availability

All data are available in the main text or the supplementary materials. The preparation of human αSyn Pre-Formed Fibrils and brain-derived αSyn fibrils of MSA- and PD-patients was conducted by Stefan Becker and Markus Zweckstetter, respectively. Human brain tissue was obtained from the Queen Square Brain Bank, UK

