## Supplementary Materials, Tables, Figures for "Human- and Rodent-derived Extracellular Vesicles Mediate the Spread of Pathology in MSA-like Models"

**Supplementary Materials and Methods**

**Proteinase-K digestion**

Enzymatic digestion of both human MSA or PD patient-derived fibrils and recombinant αSyn PFFs was performed utilizing 3 μg/mL proteinase-K for 10 min at 37°C. The enzymatic activity was terminated by addition of 1% SDS and incubation for 10 min at 65°C. Samples were then processed for immunoblotting in order to investigate the digestion patterns of total αSyn as described below.

**Trypsinization for EV surface proteins**

EV samples (0.5 mg/mL) were treated with trypsin (1 mg/mL) to eliminate the presence of loosely attached αSyn fibrils to the membranes of EV preparations. Subsequently, the reaction was terminated by addition of protease inhibitors (4 mg/mL), as previously described [52]. EVs were again isolated via centrifugation at 50.000 g (2 h, 4°C) and resuspended in PBS.

**Saponin Treatment and Measurement of EV-associated αSyn protein levels by ELISA**

EV samples were diluted in PBS to reach a final concentration of 0.7 μg/μL. 15 μL of the diluted samples were incubated with 0.5% saponin, supplemented with protease and phosphatase inhibitors at 37°C for 30 min. Following incubation, the samples were centrifuged at 50,000 g (1.5 h, 4°C). As saponine disrupts the EV membrane, the sediment corresponds to the EV membranes, whereas the supernatant includes the protein cargo present in the lumen of EVs. The latter was carefully collected for subsequent ELISA analysis to quantify αSyn levels.

**Preparation of EV-depleted cell culture medium**

EV-depleted cell culture medium was prepared following the protocol outlined by Thery et al. [50]. Essentially, DMEM supplemented with 20% FBS and 1% penicillin/streptomycin underwent overnight ultracentrifugation at 100.000 g at 4˚C using a T865 fixed-angle rotor (Sorvall Discovery). The following day, the resulting supernatant was filtered through a 0.22 μm syringe filter and stored at 4°C until needed. For all experiments, OLN-cells were cultured in EV-depleted medium containing 2% FBS.

**Micro-Bradford assay**

To determine the total protein concentration in the EV-related preparations, a standard curve was constructed using a series of BSA standards ranging in concentration from 0 to 0.6 μg/μL. Next, duplicate samples of 2 μL each were added sequentially to a flat transparent 96-well microplate (Greiner, Cat. No.: 655101), followed by incubation with 198 μL of the Bradford reagent for 5 min at RT in the dark. The absorbance was then measured at 595 nm and plotted against the concentration of BSA, allowing for the quantification of total protein in the samples.

**Concentration of EV-free Medium**

The EV-depleted supernatants were concentrated using Amicon ultra centrifugal filter units (3.000 MWCO, Cat. No.: UFC900308, Milipore). Each sample was placed in a filter unit and centrifuged at 4.000 g for 30 min at 4°C, until the final volume of the concentrated medium reached 250 μL. The concentrated medium was then collected and stored at -80°C until further analysis.

**Extracellular vesicles labeling**

Isolated EVs were labeled using the Red Fluorescent Cell Linker Kit PKH26 (Sigma Aldrich), a lipophilic dye that intercalates into the EV membrane through its long aliphatic tails and provides a stable red fluorescence signal for tracking vesicle distribution, as per the manufacturer’s instructions. Initially, the pelleted EVs were suspended in 1 mL of Diluent C, mixed with an equal volume of the dye solution containing 4×10^-6^ M PKH26 in Diluent C and incubated for 5 min at RT. The labeling reaction was terminated by adding an equal volume of 1% BSA, followed by ultracentrifugation at 100.000 g for 2 h at 4°C to obtain the pellet that contained the PKH26-labeled EVs.

**Behavioral tests**

Motor coordination and balance were assessed using the rotarod performance test and the beam traversal test, following previously published protocols [53, 54]. For the rotarod test, mice were placed on a rotating rod with gradually accelerating speed. Each mouse underwent three trials with an inter-trial interval of 120 min. A trial concluded when the animal either fell off the rod or grabbed onto the apparatus, rotating around it twice in succession without attempting to move across the rod. Latency to fall (seconds) was recorded for each trial and the average latency was used for statistical analysis. For the beam traversal test, mice were trained to traverse a gradually narrowing beam elevated above a surface over two days prior to testing. On the testing day, the beam was covered with a metallic grid and each mouse was placed at the widest end of the beam and allowed to walk to a safety platform at the opposite, narrower end. The trials concluded when the mouse inserted one of its forelimbs into the cage. Mice underwent five trials. Performance was assessed by measuring the number of foot slips during forward motion and the time taken to cross the beam. The averages of these parameters were used for statistical analysis.

**DAB staining**

The brain sections were incubated for 10 min in a mixture of 3% H_2_O_2_ and 10% methanol. Next, a 5% NGS blocking solution was added to minimize non-specific antibody binding. The primary antibody (anti-tyrosine hydroxylase (TH; 1:5.000) or anti-pSer129-αSyn (EP1536Y; 1:10.000)) was applied for 24 h, followed by 1 hour incubation with a secondary biotinylated anti-rabbit antibody (1:3000; Cat.No.: PK-4001, Vector Laboratories) in 2% NGS. An avidin-biotin peroxidase complex system (ABC Elite; Cat.No.: PK-4001, Vector Laboratories) was then utilised for 1 hour at RT to enhance the detection of the immunocomplexes, which were visualized using the chromogen 3,3′-diaminobenzidine (DAB; Cat.No.: K3468, Dako). Subsequently, the sections were dehydrated through a series of graded alcohols and coverslipped. For TH-staining, the sections from the striatum were digitally scanned using a Prime Histo XE slide scanner to evaluate the TH^+^ staining, which appeared as a brown precipitate, whereas for pSer129-αSyn staining brightfield microscopy was utilized to analyse sections from olfactory bulbs, prefrontal cortex, striatum, hippocampus, substantia nigra and cerebellum.

**Supplementary Tables**

| ***Table S1***: **Demographics of donor brains from MSA and PD patients from where the MSA and PD brain-amplified fibrils were generated** | | | | | |
| --- | --- | --- | --- | --- | --- |
| **Disease type** | **Age** | **Gender** | **Postmortem delay (hours)** | **Cause of death** | **Disease duration (years)** |
| MSA1 | 82 | Male | 8 | Cardiorespiratory failure | 7 |
| MSA2 | 71 | Female | 19 | Hypostatic pneumonia | 6 |
| PD | 79 | Male | 17 | Acute myocardial infarction | 7 |

| ***Table S2***: **Demographics of donor brains from MSA patients and controls obtained from the Queen Square Brain Bank** | | | | | |
| --- | --- | --- | --- | --- | --- |
| **Case number** | **Age** | **Gender** | **Postmortem delay (hours)** | **Pathological diagnosis** | **Disease duration (years)** |
| CTR 1 | 103 | Female | 26 | Control Brain, Intermediate level Alzheimer’s disease neuropathological change , Cerebral amyloid angiopathy, Atheroma of leptomeningeal vessels. | - |
| CTR 2 | 96 | Male | 38 | Control Brain, Pathological ageing, Argyrophilic grain disease, Cerebral amyloid angiopathy | - |
| MSA 1 | 63 | Female | 46 | MSA, involving olivopontocerebellar and to a lesser extent striatonigral regions, Pathological ageing | 14 |
| MSA 2 | 65 | Female | 53 | MSA-C (OPCA) | 10 |

| ***Table S3*: List depicting the primary and secondary antibodies, as well as dyes used in Western blotting (WB), immunocytochemistry (ICC) or immunohistochemistry (IHC)** | | | | | | | | | | |
| --- | --- | --- | --- | --- | --- | --- | --- | --- | --- | --- |
| **Primary Antibodies against** | **Clone** | **Cat. No** | **Working dilutions**  **WB ICC IHC** | | | | | | **Species reactivity** | **Company** |
| Human αSyn | 4B12 | GTX21904 | 1:1000 | | - | | - | | mouse | Gene Tex |
|  | LB-509 | 807701 | - | | 1:1000 | | - | | mouse | Biolegend |
| Total αSyn | D10 | Sc-515879 | - | | 1:1000 | | - | | mouse | Santa Cruz |
|  | Syn1 | 610787 | 1:1000 | | - | | - | | mouse | BD Transductions |
| Rodent αSyn | D37A6 | 4179 | 1:1000 | | 1:400 | | - | | rabbit | Cell Signaling |
| pSer129-αSyn | EP1536Y | ab51253 | 1:1000 | | 1:100 | | 1:1000 | | rabbit | Abcam |
| Oxidized/ Nitrated αSyn | SYN303 | 824301 | - | | 1:1000 | | - | | mouse | Biolegend |
| Aggregated αSyn | MJFR-  14-6-4-2 | ab209538 | - | | 1:5000 | | - | | rabbit | Abcam |
| TPPP/p25α |  | gift | 1:1000 | | - | | - | | rabbit | Home-made |
| β-actin |  | TA309077 | 1:2000 | | - | | - | | mouse | OriGene |
| Alix |  | 2171 | 1:1000 | | - | | - | | mouse | Cell Signaling |
| Flotillin-1 |  | sc-133153 | 1:1000 | | - | | - | | mouse | Santa Cruz |
| TSG101 |  | Ab125011 | 1:1000 | | - | | - | | rabbit | Abcam |
| CD9 |  | ab307085 | 1:1000 | | - | | - | | rabbit | Abcam |
| BSA |  | #ΗΥΒ 267-01 | 1:1000 | | - | | - | | mouse | ANTIBODY SHOP, BioPorto |
| TUJ1 |  | MMS-435P | - | | 1:2000 | | - | | mouse | Bioline Covance |
| MAP2 |  | sc-20172 | - | | 1:500 | | - | | rabbit | Santa Cruz |
| MBP |  | MSA409S | - | | 1:200 | | 1:200 | | rat | Biorad |
| TH |  | AB76442 | - | | - | | 1:2000 | | chicken | Abcam |
| **Secondary Antibodies** | **Cat. No.** | **Working dilutions**  **WB ICC IHC** | | | | | | | **Species**  **reactivity** | **Company** |
| Goat Anti-Mouse gG- HRP  conjugated | AP124P | 1:5000 | | - | | - | | | mouse | Sigma Aldrich |
| Goat Anti-Rabbit IgG-HRP  conjugated | AP132P | 1:5000 | | - | | - | | | rabbit | Sigma Aldrich |
| CF555 red | 20033 | - | | 1:2000 | | 1:2000 | | | rabbit | Biotium |
|  | 20096 | - | | 1:2000 | | - | | | rat | Biotium |
| CF488A green | 20012 | - | | 1:2000 | | - | | | rabbit | Biotium |
|  | 20010 | - | | 1:2000 | | - | | | mouse | Biotium |
|  | ab13970 | - | | - | | 1:2000 | | | chicken | Abcam |
| Cy5 | 115-175-146 | - | | 1:500 | | 1:500 | | | mouse | Jackson Imm. Affinipure |
|  | 712-175-153 | - | | - | | 1:500 | | | rat | Jackson Imm. Affinipure |
| **Dyes** | **Cat. No.** | **Working dilutions**  **WB ICC IHC** | | | | | | | | **Company** |
| DAPI | 10236276001 | - | | 1:3000 | | | | 1:3000 | | Sigma Aldrich |
| General Cell Membrane Labeling | PKH26GL | - | | 2x10^–6^ M | | | | 2x10^–6^ M | | Sigma Aldrich |

**Supplementary Figure Legends**

**Figure S1:** **Addition of MSA1, MSA2 or PD patient-amplified fibrils to rat oligodendroglial cell lines is accompanied by the formation of Triton-, SDS- and UREA-soluble αSyn species.**

(A) Representative transmission electron microscopy images of negatively stained MSA1, MSA2 or PD patient-derived fibrils, demonstrating the ultrastructural architecture of the different fibril types. Scale bar: 500 nm. (B) Representative immunoblot of total αSyn (Syn1), demonstrating the distinct digestion profiles of αSyn species generated upon digestion with proteinase K (3 μg/mL) of PFFs, MSA1, MSA2 or PD patient-amplified fibrils (1.5 μg/mL). (C-E) Representative immunoblots of human (4B12) (top row) and total (Syn1) (lower row) αSyn protein levels in the Triton-, SDS- and UREA-soluble fractions of OLN cells treated with 1 μg/mL MSA1 (C), MSA2 (D) or PD (E) fibrils (or PBS as control) for 48h or 8 days. (F-H) Quantifications of the intracellular (Triton-, SDS- and UREA-soluble) human and total αSyn protein levels. β-actin detection confirms equal loading. Data are expressed as the mean ± SE of at least three independent experiments; *p<0.05; **p<0.01; ***p<0.001, by one-way ANOVA with Tukey’s post hoc test, comparing PBS- and fibril-treated conditions within the same cell line and #p<0.05; ##p<0.01; ###p<0.001, by two-way ANOVA with Bonferroni’s correction, comparing between the different OLN cells.

**Figure S2:** **Addition of high amounts of human MSA1, MSA2 or PD patient-amplified fibrils to rat oligodendroglial cell lines augments the formation of Triton-insoluble αSyn species.**

(A-C) Representative immunoblots of human (4B12) (top row) and total (Syn1) (lower row) αSyn protein levels in the Triton-, SDS- and UREA-soluble fractions of OLN cells treated with 6 μg/mL MSA1 (A) MSA2 (B), or PD (C) fibrils (or PBS as control) for 48h or 8 days. (D-F) Quantifications of the intracellular (Triton-, SDS- and UREA-soluble) human and total αSyn protein levels. β-actin detection confirms equal loading. Data are expressed as the mean ± SE of at least three independent experiments; *p<0.05; **p<0.01; ***p<0.001, by one-way ANOVA with Tukey’s post hoc test and #p<0.05; ##p<0.01; ###p<0.001, by two-way ANOVA with Bonferroni’s correction.

**Figure S3: Human MSA or PD patient-derived fibrils evoke the recruitment of the endogenous rodent oligodendroglial αSyn and its incorporation into pathological αSyn assemblies.**

(A-C) Representative immunofluorescence images using antibodies against endogenous rodent (A-B, red, D37A6), human (A, green, LB-509), oxidized/nitrated (B, green, SYN303), aggregated (C, red, MJFR-14-6-4-2) and total (C, green, D10) αSyn and DAPI staining in OLN cell lines treated with PBS or 1 μg/mL of MSA1, MSA2 or PD fibrils for 48h or 8 days. Scale bar: 25 μm. (D-H) Quantifications of endogenous rodent (D), human (E), oxidized/nitrated (F), aggregated (G) and total (H) protein levels in OLN-93, OLN-AS7 and OLN-p25α cells measured as area surface/cell following treatment with 1 μg/mL patient-derived fibrils for 48h or 8 days. Data are expressed as the mean ± SE of three independent experiments with triplicate samples per condition within each experiment; *p<0.05; **p<0.01; ***p<0.001, by one-way ANOVA with Tukey’s post-hoc-test and #p<0.05; ##p<0.01; ###p<0.001, by two-way ANOVA with Bonferroni’s correction.

**Figure S4: Incubation of OLN cells with high amounts of human patient-derived fibrils evokes the formation of aberrant αSyn conformations incorporating the endogenous oligodendroglial protein that decrease over time.**

(A-D) Representative immunofluorescence images using antibodies against endogenous rodent (A-B, red, D37A6), human (A, green, LB-509), oxidized/nitrated (B, green, SYN303), aggregated (C, red, MJFR-14-6-4-2), total (C-D, green, D10) and pSer129-αSyn (D, red, EP1536Y) αSyn and DAPI staining in OLN cell lines treated with 6 μg/mL of MSA1, MSA2 or PD fibrils for 48h or 8 days. Scale bar: 25 μm. (E-J) Quantifications of endogenous rodent (E), human (F), oxidized/nitrated (G), aggregated (H), total (I) and pSer129-αSyn (J) protein levels in OLN-93, OLN-AS7 and OLN-p25α cells measured as area surface/cell following treatment with 6 μg/mL patient-derived fibrils for 48h or 8 days. Data are expressed as the mean ± SE of three independent experiments with triplicate samples per condition within each experiment; *p<0.05; **p<0.01; ***p<0.001, by one-way ANOVA with Tukey’s post-hoc-test and #p<0.05; ##p<0.01, by two-way ANOVA with Bonferroni’s correction.

**Figure S5:** **Treatment of rat oligodendroglial cells with** **human MSA2 patient-derived fibrils enhances EV-associated αSyn release.**

(A) Graphs depicting the concentration and size distribution of EVs released from oligodendrocytes treated with ΜSA1, MSA2, PD fibrils or PFFs (1 or 6 μg/mL, 48h or 8 days) (or PBS). (B-C) Immunoblots demonstrating enhanced human and total αSyn secretion via ΕVs from OLN cells treated with 1 (B) or 6 μg/mL (C) MSA2 fibrils for 48h or 8 days. (D-E) Quantifications of human (top) and total (bottom) αSyn in EV-related fraction of OLN cells treated with 1 (D) or 6 μg/mL (E) MSA2 fibrils for 48h or 8 days. Alix confirms equal loading. Data are expressed as the mean ± SE of at least three independent experiments; *p<0.05; **p<0.01; ***p<0.001, by one-way ANOVA with Tukey’s post-hoc-test and #p<0.05; ##p<0.01, by two-way ANOVA with Bonferroni’s correction. (F) Immunoblots of human (4B12) and total (Syn1) αSyn protein levels of trypsinized or non-trypsinized EVs, isolated from OLN cells incubated with 1 μg/mL PD fibrils for 48h (or PBS). Alix verifies equal loading. (G) Quantification of total αSyn in saponine-treated EVs from OLN-AS7 cells treated with 1 μg/mL PD fibrils for 48h or 8 days (or PBS).

**Figure S6: Human MSA type 2 patient-derived fibrils induce the phosphorylation of the endogenous oligodendroglial αSyn at Ser129 and accelerate its release via EVs.**

(A-B) Representative immunoblots of rodent and pSer129-αSyn protein levels (D37A6 and EP1536Y antibodies, respectively) in the Urea-soluble (A) and EV-associated fractions (B) of OLN-93, OLN-AS7 and OLN-p25α cells treated with 6 μg/mL MSA type 2 fibrils (or PBS as control) for 48h or 8 days. (C-D) Quantifications of the Urea-soluble (C) and EV-associated (D) rodent (left panels) and pSer129-αSyn (right panels) protein levels. Equal loading was verified by the detection of β-actin (total protein marker) and alix (EV marker). Data are expressed as the mean ± SE of three independent experiments; *p<0.05; **p<0.01; ***p<0.001, by one-way ANOVA with Tukey’s post-hoc-test and #p<0.05; ##p<0.01, by two-way ANOVA with Bonferroni’s correction.

**Figure S7: Effects of human MSA2 patient-derived fibrils on intracellular and extracellular TPPP/p25α levels over time.**

(A-B) Representative immunoblots (upper panels) and quantifications (bottom panels) of Triton-, SDS-, Urea-soluble and EV-associated TPPP/p25α protein levels in OLN-p25α cells incubated with 1 (Α) or 6 μg/mL (B) human MSA type 2 fibrils (or PBS as control) for 48h or 8 days. β-actin and alix were used as loading controls. Data are expressed as the mean ± SE of three independent experiments; *p<0.05; **p<0.01, by one-way ANOVA with Tukey’s post-hoc-test. (C) Representative immunoblots of extracellular EV-free TPPP/p25α protein levels in OLN-p25α cells treated with 6 μg/mL human MSA type 2 (or PBS as control) for 48h or 8 days. Equal loading was verified by the detection of BSA. Data are expressed as the mean ± SE of three independent experiments. (D) Representative immunofluorescence images using antibodies against TPPP/p25α (grey, p25α), endogenous rat αSyn (red, D37A6), human αSyn (green, LB-509) and DAPI staining in OLN-p25α cells treated with PBS, 1 μg/mL MSA type 1 or PD fibrils for 48h. Scale bar: 25 μm.

**Figure S8: MSA2-fibril-treated OLN cell-derived EVs are taken up by primary murine cortical neurons and induce the formation of pathological αSyn assemblies.**

(A) Immunofluorescence images (left) and 3D-reconstructed image (right), depicting the uptake of PHK26-labeled OLN-p25α derived EVs uptake (40 µg/mL) by mouse primary cortical neurons (TUJ1^+^) following overnight incubation. Scale bar: 25 µm. (B-F) Representative immunofluorescence images using antibodies against (B) neuronal dendrites (red, MAP2), human αSyn (green, LB-509), (C) neuronal axons (green, TUJ1), rodent αSyn (red, D37A6), (D) neuronal dendrites, oxidized/nitrated αSyn (green, SYN303), (E) neuronal axons, aggregated αSyn (red, MJFR-14-6-4-2), (F) neuronal axons, pSer129-αSyn (red, EP1536Y) and DAPI staining in mouse primary neurons inoculated with OLN cell-derived EVs (40 µg/mL for 72 h or 6 days in F), previously treated with 6 µg/mL MSA2 fibrils for 48h. Scale bar: 25 µm. (G) Quantifications of human, rodent, oxidized/nitrated, aggregated and pSer129-αSyn protein levels expressed as area surface/cell. Data are expressed as the mean ± SE of three independent experiments with triplicate samples/condition; *p<0.05; **p<0.01; ***p<0.001, by one-way ANOVA with Tukey’s post-hoc-test and ###p<0.001, by two-way ANOVA with Bonferroni’s correction.

**Figure S9: The levels of seeded pathological αSyn conformations engendered within mouse primary cultures upon addition of fibril-treated oligodendroglial EVs depend on the endogenous αSyn expression. Both TUJ1^+^ axonal and MAP2^+^ dendrite networks remain relatively unaffected.**

(A-C) Immunofluorescence images using antibodies against (A) neuronal axons (green, TUJ1), rodent αSyn (red, D37A6), (B) neuronal dendrites (red, MAP2), human αSyn (green, LB-509), (C) neuronal dendrites, oxidized/nitrated αSyn (green, SYN303) and DAPI staining in KO-αSyn mouse cortical neurons inoculated with EVs (40 µg/mL for 72 h) from OLN cells treated with 6 µg/mL PD fibrils for 48h (or PBS). (D-G) Quantifications of TUJ1 (D-E) and MAP2 (F-G) protein levels in mouse primary cortical cultures measured as area surface/cell following incubation with EVs (40 µg/mL for 72 h) derived from OLN cells previously treated with 6 µg/mL MSA1 or PD patient-derived fibrils or PFFs (D.F) or with 6 µg/mL MSA2 (E,G) for 48h (or PBS as control). Data are expressed as the mean ± SE of three independent experiments with triplicate samples/condition within each experiment.

**Figure S10: EVs from OLN cells incubated with human MSA2 patient-derived fibrils seed the templating of the endogenous oligodendroglial αSyn into the formation of pathological αSyn species within rat primary oligodendroglial cultures.**

(A) Immunofluorescence images (left) and 3D-reconstructed image (right), depicting the uptake of PKH26-labeled oligodendroglial-derived EVs (40 µg/mL) by mature rat oligodendrocytes (p25α^+^) following overnight incubation. DAPI marked nuclei. Scale bar: 25 µm. (B-D) Representative immunofluorescence images using antibodies against (B) oligodendrocytes (red, MBP), rodent αSyn (green, D37A6), oxidized/nitrated αSyn (grey, SYN303), (C) oligodendrocytes, aggregated αSyn (green, MJFR-14-6-4-2), human αSyn (grey, LB-509), (D) oligodendrocytes, pSer129-αSyn (green, EP1536Y), total αSyn (grey, D10) and DAPI staining in rat oligodendrocytes inoculated with EVs (40 µg/mL for 72 h or 6 days in D) from OLN cells pre-treated with 6 µg/mL MSA2 fibrils for 48h (or PBS). Scale bar: 25 µm. (E) Quantifications of rodent, oxidized/nitrated, aggregated, human, pSer129, total αSyn and MBP protein levels in rat oligodendrocytes measured as area surface/cell following incubation with OLN-derived EVs. Data are expressed as the mean ± SE of three independent experiments with triplicate samples/condition; *p<0.05; **p<0.01; ***p<0.001, by one-way ANOVA with Tukey’s post-hoc-test and #p<0.05; ###p<0.001, by two-way ANOVA with Bonferroni’s correction.

**Figure S11: Widespread pSer129-αSyn protein expression across the rostrocaudal axis of the PLP-hαSyn mouse brain, as compared to WT-αSyn brain.**

Representative brightfield microscopy images of DAB-stained brain sections derived from six-month-old WT-αSyn and PLP-hαSyn mice using a primary antibody against pSer129-αSyn (EP1536Y; 1:10.000), revealing enhanced immunoreactivity for pSer129-αSyn^+^ signal in various brain regions of PLP-hαSyn mice, as compared to the respective ones of WT-αSyn mice. Brain regions examined include the olfactory bulb, prefrontal cortex, striatum, hippocampus, substantia nigra and cerebellum. Lower (x20, left) and higher (x63, right) magnification images of different areas (indicated by the arrows) for each region/genotype are shown. Scale bar: 25 μm.

**Figure S12: Intrastriatal delivery of WT-αSyn or PLP-hαSyn-derived EVs does not alter the integrity of the nigrostriatal axis of WT-αSyn mice, assessed at one and three months post-injection.**

(A) Representative TH-DAB immunostaining in coronal striatal sections of WT-αSyn mice inoculated with WT-αSyn or PLP-hαSyn EVs, assessed at one (upper) or three (bottom) months post-EV injection. (B) Quantifications of dopaminergic fibre density relative to the corresponding contralateral hemisphere in WT-αSyn striata injected with WT-αSyn or PLP-hαSyn brain-derived EVs at one and three months post-injection (six sections per animal, with a 5-section interval). Data are expressed as the mean ± SE (n=6 animals per group). (C) Representative TH immunofluorescence of ventral midbrain sections derived from WT-αSyn mice at one (upper) or three (bottom) months post-injection with with EVs isolated from WT-αSyn or PLP-hαSyn mouse brains. Scale bar: 100 µm. (D) Measurement of the total TH^+^ neurons in the contralateral and ipsilateral SNpc of WT-αSyn mice injected with EVs isolated from WT-αSyn or PLP-hαSyn mouse brains at one and three months post-injection (ten sections per animal, with a 3-section interval). Data are expressed as the mean ± SE (n=6 animals per group).

**Figure S13: Intrastriatal administration of human MSA-treated OLN-p25α-derived EVs does not affect dopaminergic neuron survival or motor performance and coordination of PLP-hαSyn mice, at three months post-delivery.**

(A) Representative TH-DAB immunostaining in coronal striatal sections of PLP-hαSyn mice unilaterally injected with OLN-p25α-derived EVs, pre-incubated with PBS or MSA patient-derived fibrils, three months post-inoculation. (B) Quantifications of dopaminergic TH fibre density normalized to the corresponding contralateral hemisphere of PLP-hαSyn mouse striatum did not reveal significant differences between groups. Data are expressed as the mean ± SE (n=3 mice/group). (C) Representative TH immunofluorescence images of PLP-hαSyn mouse ventral midbrain sections, assessed at three months post-intrastriatal delivery of EVs isolated from OLN-p25α cells treated with PBS- or human MSA patient-amplified αSyn fibrils. Scale bar: 100 µm. (D) Quantifications of the total TH^+^ neuron cell number in the SNpc of PLP-hαSyn mice did not reveal significant changes between groups. Data are expressed as the mean ± SE (n=3 mice/group). (E) Behavioral assessments, including the challenging beam test (errors per step and traversal time) and rotarod test (latency to fall), demonstrated no significant differences in motor performance or coordination amongst groups. Data are presented as the mean ± SE (n=3 mice/group).

**Figure S14: Inoculation of brain-derived EVs isolated from MSA patients does not affect dopaminergic system integrity or motor phenotype of recipient PLP-hαSyn mice.**

(A) Representative TH-DAB immunostaining in coronal striatal sections of PLP-hαSyn animals injected with EVs originating from human MSA or CTR brains, at three-months post-inoculation. (B) Quantification of dopaminergic TH fibre density in proportion to the corresponding contralateral hemisphere in the striatum of PLP-hαSyn mice injected with human MSA or CTR brain-derived EVs. Data are expressed as the mean ± SE (n=4 mice/group). (C) Representative TH immunofluorescence ventral midbrain images of PLP-hαSyn mice inoculated with EVs from human MSA or CTR brains, at three-months post-inoculation. Scale bar: 100 µm. (D) Quantifications of the total nigral TH^+^ neuron number of PLP-hαSyn mice injected with human MSA or CTR brain-derived EVs. Data are expressed as the mean ± SE (n=4 mice/group). (E) Behavioral assessments, including the challenging beam test (errors per step and traversal time) and rotarod test (latency to fall), did not uncover significant differences in motor performance or coordination among groups. Data are expressed as the mean ± SE (n=4 mice/group).
